# Solvent-induced allosteric transition of the Hepatitis C virus human cellular receptor CD81 large extracellular loop

**DOI:** 10.1101/2022.03.21.485124

**Authors:** C. Risueño, D. Charro, N. G. A. Abrescia, I. Coluzza

**Affiliations:** Structure and Cell Biology of Viruses Lab, Center for Cooperative Research in Biosciences (CIC bioGUNE), Basque Research and Technology Alliance (BRTA), Derio Spain; Center for Cooperative Research in Biomaterials (CIC biomaGUNE), BRTA, San Sebastian, Spain; BCMaterials, Basque Center for Materials, Applications and Nanostructures, Bld. Martina Casiano, UPV/EHU Science Park, Barrio Sarriena s/n, 48940 Leioa, Spain; IKERBASQUE, Basque Foundation for Science, Bilbao Spain; Centro de Investigación Biomédica en Red de Enfermedades Hepáticas y Digestivas (CIBERehd), Instituto de Salud Carlos III, Madrid Spain

**Keywords:** tetraspanin CD81, Hepatitis C virus, cell-entry, molecular dynamics, cell signalling

## Abstract

CD81 is a tetraspanin receptor that clusters into microdomains to mediate cell signalling processes. CD81 is also one of the four primary cellular receptors of the Hepatitis C virus (HCV). Previous structural studies on the α-helical CD81 large-extracellular-loop domain (CD81_LEL_) have shown that it can adopt different conformations (from closed to open), likely depending on the environmental conditions. This conformational plasticity has been implicated in the endosomal fusion of HCV upon entry. However, the precise mechanism governing the CD81_LEL_ plasticity has remained elusive so far.

Here, by combining molecular dynamics simulations and circular dichroism experiments on wt-CD81_LEL_ and two mutants at different endosomal pH conditions, pH 5.5 and pH 4.6, we show that the modulation of the solvation shell governs the plasticity of CD81_LEL_. The primarily implicated residues are D139 and E188, respectively, located near a loop preceded by a helix. At acidic conditions, their interaction with water is reduced, causing a re-ordering of the water molecules, and thus triggering the dynamics of CD81_LEL_. However, mutations E188Q and D139N retain the solvation shell and restrict the conformational space that the head subdomain can explore.

We propose that residues E188 and D139 control the solvent-induced allosteric transition of the CD81_LEL_ domain. This mechanism might play a role in other cellular receptors that function along the endosomal pathway.

**Popular Summary:** Understanding the cellular mechanisms that are exploited by viruses to infect their host is key for the development of therapeutics. Here, in the context of Hepatitis C Virus infection we report the mechanism that governs the plasticity of the extra-cellular domain of tetraspanin CD81, one of the major cellular receptors of this virus. The mechanism proposed here is a novel form of solvent-induced allosteric transition in proteins mediated by two antenna residues located in the head subdomain of CD81. We propose that it could serve as a pH sensing strategy to time the endosomal pathway and trigger a signal at the right time for HCV fusion.

## I. Introduction

Tetraspanins are a family of cell-surface membrane proteins characterised by four transmembrane domains, one small extracellular loop (SEL) and a large extracellular loop (LEL). When a partner protein approaches, tetraspanins cluster, forming tetraspanin-enriched microdomains (TEMs) to amplify and transduce signals that regulate a wide variety of cellular functions. Cluster of Differentiation 81 (CD81) is a human cell surface tetraspanin, that apart from if physiological signalling activities, has also been shown to participate in tumorigenesis, cancer progression and infectious processes [1–3]. The CD81 protein, among a few other tetraspanins, is considered a classical marker for exosomes, their biogenesis and their involvement in vesicle trafficking [4]. Vesicle trafficking, protein sorting, and targeted degradation of some sorted cargo are biological processes strictly linked to the gradual maturation of endosomes into a lysosome and acidification of the late endosome [5]. CD81 is also one of the primary cellular receptors for the entry of the Hepatitis C virus (HCV) [1]. HCV is a pleomorphic single-stranded positive RNA enveloped virus (family *Flaviviridae*) with glycoproteins E1 and E2 decorating the viral envelope. HCV virus cell entry is a complex process involving several host factors [6]. Early cellular studies at pH 7 and pH 5 on culture-produced HCV particles have shown that HCV entry is pH-dependent [7]. Later studies have demonstrated that after clathrin-mediated endocytosis of the virus and delivery to early endosomes, it remains in the endosome until an internal pH is achieved, which stimulates fusion between the viral envelope and the endosomal membrane [8,9]. Acidification of endosomes is critical for viral fusion activity. Different enveloped viruses show distinct pH triggering values (*i*.*e*. Semliki-Forest virus requires pH 6.1 or below, Influenza ranges from pH 5.3 to 5.9 depending on the strain, Dengue 2 virus requires pH 5.5, etc.; while in general, *Flaviviruses* require a pH between pH 6.4 and 4.8) [10]. While early endosomes maintain a pH of about 6.8-6.1, late endosomes of a pH from 6.0 to 4.8 and lysosomes can drop to pH values of 4.5 [11]. Biochemical and cellular experiments indicate that CD81 primes conformational changes on the E2 glycoprotein for virus’s fusion at low pH [12].

The structure of the full-length CD81 and of the large extracellular loop domain (CD81_LEL_) has been previously determined [13]. The CD81 crystal structures show five helices composing the LEL region, two forms the stalk subdomain (A and E) and three the head subdomain (B, C and D), the SEL region is disordered, and four transmembrane helices (TM 1-4) (Fig. 1A). Helices C and D are responsible for the interaction with the viral E2 [14,15]. Early structural analyses have shown that helices C and D in the head subdomain of CD81_LEL_ can adopt different conformations [16,17]. A more recent study has been able to classify three distinct CD81_LEL_ conformations, open, intermediate and close, and to relate the adopted conformation to the environmental pH and redox conditions [18]. This led to the proposal that the conformational dependence of the human CD81_LEL_ from pH and redox conditions enabled the virus-receptor interactions to re-engage at endosomal conditions [18]. However, the precise molecular mechanisms that would make the CD81_LEL_ structure respond to changes in pH has remained elusive so far.

**FIG 1.**
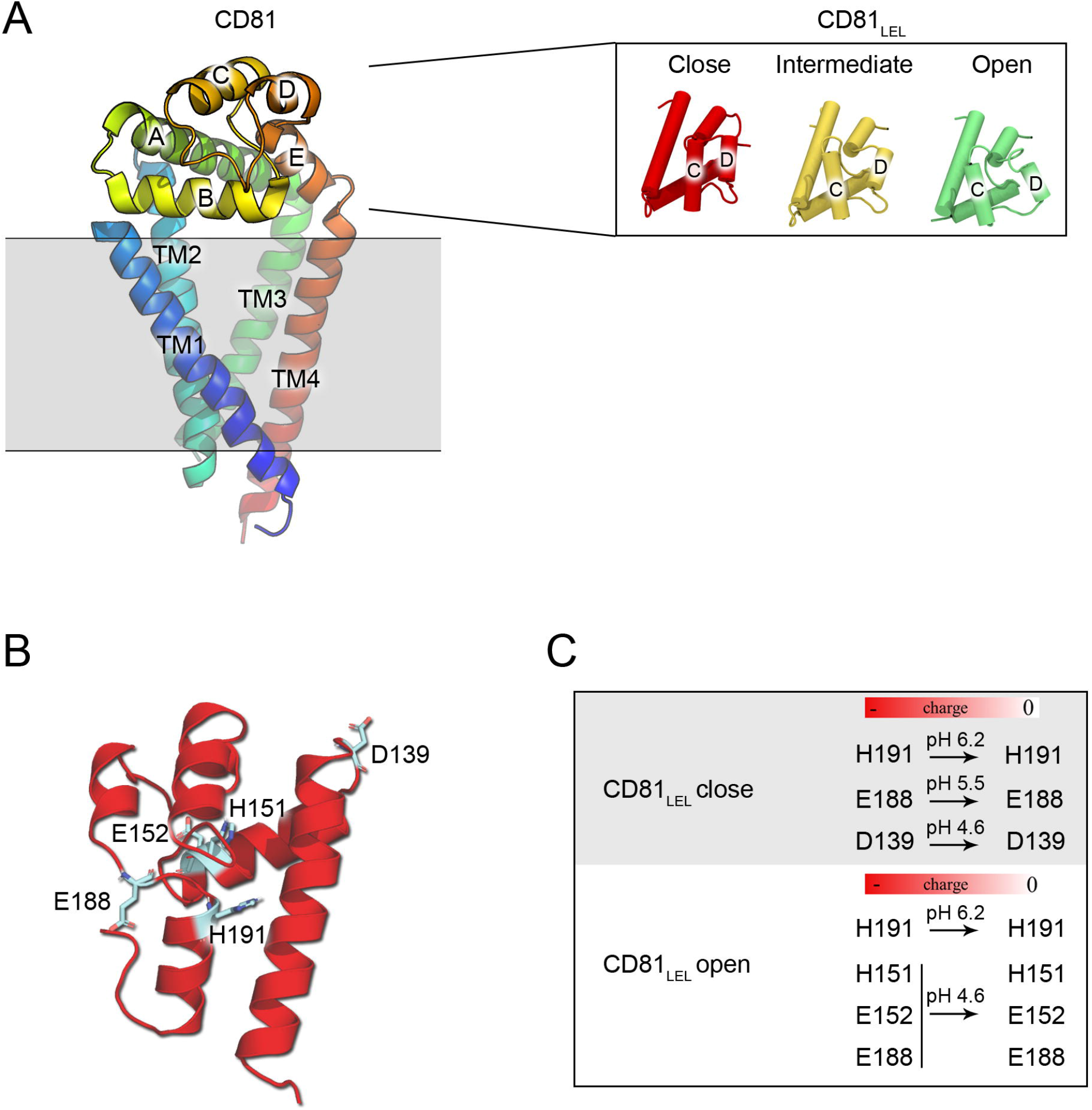
Structure of CD81. (A) Crystal structure of the full-length CD81 represented in cartoon and colour-coded from blue to red from the N-terminus to the C-terminus, adapted from. Inset, the three different crystallographic conformers displayed by the CD81_LEL_ domain, adapted from. (B) Structure of the CD81_LEL_ in close conformation represented in cartoon with pH-dependent residues depicted in sticks (PDB ID 5M3T). (C) pH-dependent residues of the different CD81_LEL_ conformers at low pH.

Here, we have performed molecular dynamics (MD) simulations on CD81_LEL_ along with the pH range of the endosomal pathway and found that titration of specific residues, E188 and D139, at low pH disrupts the head subdomain’s solvation profile of the close structure, triggering the free movement of helices C and D to adopt a relaxed/open disposition. In-silico mutations at those positions, E188Q and D139N, block the influence of pH over the head subdomain’s solvation shell, restricting the conformational space that helices C and D could explore. These observations were tested through thermal stability analysis by circular dichroism (CD) of the mutated recombinantly expressed proteins.

All together our findings indicate that the liquid microenvironment around the CD81_LEL_ is an active player for the plasticity of the head subdomain. Residues D139 and E188 act as pH sensors to trigger the conformational change by modulating the solvent around the head subdomain. Once internalised and still bound to CD81, HCV could exploit such mechanism to sense and set the time for fusion. The CD81 reactivity to environmental pH might therefore be a general attribute of the molecule to discern biological cues along the endosomal pathway such as protein recycling or lysosomal escape.

## II. Material and methods

### A. In silico titration of the CD81_LEL_ crystal structures

The atomic models of the different conformations of CD81_LEL_ (residues 112-204): open (PDB ID 5M33, chain B) and close (PDB ID 5M3T, chain B); were prepared for structural modelling with the PDB2PQR server using AMBER forcefield at different endosomal pH conditions [18,19]. The pKa of the pH sensitive residues of the protein is calculated with the PDB2PQR web server which, at a desired pH, calculates the pKa that each HIS, ASP and GLU residue of a protein would have based on hydrogen bonding, local energetics, and model pKa values [19]. PropKa tool was also used for this purpose [20]. State of the art estimation of the protonation state of the residue uses classical approaches with remarkable accurate results [20–23]. Hence, quantum effects were not considered and are probably beyond the scope of this work. The changes in the electrostatic potential of CD81_LEL_ at different endosomal pH values (from 7.4 to 4.6) of the resulting structures were then analysed with the APBS tool interfaced within the molecular visualization package VMD [24].

### B. Molecular dynamics

All MD simulations were performed using AMBER99SB-ILDN force field and explicit water solvent (TIP4P model) with GROMACS software 2019.1 version [25]. After structures setup (hydrated and neutralised) in a cubic box size of 10 nm, an equilibration step was applied to the resulting systems, allowing them to relax for 500 ps, using V-rescale and Berendsen algorithms for the temperature and pressure equilibration, respectively. These equilibrated structures were then used for the production trajectories. Trajectories were performed at constant pressure (1 bar) and physiological temperature (309 K) using standard coupling schemes, Parrinello-Rahman for the pressure coupling and Nose-Hoover for the temperature.

We simulated our system in five different pH conditions described as characteristic of the endosomal pathway: 7.4, 6.2, 5.5, 5.0 and 4.6. The pH was set up in the topology of the system, by selecting in the interactive interface of GROMACS which residues (LYS, ARG, GLU, ASP, HIS, GLN) of the protein should be protonated or not. This was done after the PBD2PQR analysis, in which we inspected the pKa values of the protein at a desired pH. We then regularly checked along the trajectory for changes in protonation state of the residues. The PBD2PQR has shown to be an effective tool in predicting the residue pKa value for a static conformation. We then coupled the protonation state of the residue to the MD in a self-consistent scheme. If we found alterations, we recomputed the topology of the system and restarted the simulation with the new protonation. We found that our system quickly converged to a particular protonation state and did not change anymore. The efficiency of our constant pH scheme is admittedly not general because it depends on the fluctuations in the protonation state peculiar to each system. However, it was relatively simple to implement and suited our needs. Moreover, although several strategies have been proposed it is only recently that GROMACS has implemented the λ-dynamics methods in the main software package [21–23]. We also studied the effect of artificially fixing the titratable state of CD81_LEL_ close into the open conformation and vice versa on the system’s enthalpy (see supplemental material Text-S1). Two sets of MD experiments were performed: (i) 2 ns simulations to study the evolution of the enthalpy of CD81_LEL_ open and close at different endosomal pH; (ii) 1μs simulations to study the plasticity of CD81_LEL_ at different endosomal pH.

Fifty-two MD simulations were performed in the i2BASQUE supercomputing network, the ATLAS machine at the DIPC and the Spanish Supercomputing Network (RES) facilities. The resulting trajectories were analysed using VMD and the Root Mean Square Deviation (RMSD) trajectory tool, which shows differences between two structures by calculating the difference in position of the atoms of the -new-structure as a function of time, with respect to the reference structure. Protein-solvent interactions were analysed using the Radial Pair Distribution g(r) function which consists on calculating the density of particles surrounding a reference particle within an radius r + dr (inner and outer radius, respectively) [24,26].

For each residue and pH we computed the separate contributions to the protein enthalpy H^P^_pH_ = H^PP^_pH_ + H^PS^_pH_, namely the intra-protein interactions (H^PP^_pH_) and the residue-solvent interactions (H^PS^_pH_) averaged over water conformations generated through short trajectories. We then consider the system at pH 7.4 as a reference and compute the differences ΔH^PP^_pH_ = H^PP^_pH_ − H^PP^_pH 7.4_ and ΔH^PS^_pH_ = H^PS^_pH_ − H^PS^_pH 7.4_. Finally, we calculated the differences between the open and close conformers, ΔΔH^PP^_pH_ = ΔH^PP^_pH,Open_ − ΔH^PP^_pH,Close_ and ΔΔH^PS^ = ΔH^PS^_pH,Open_ − ΔH^PS^_pH,Close_.

### C. In silico mutation of the CD81_LEL_ crystal structures

In silico mutations of CD81_LEL_ in open (PDB ID 5M33, chain B) or close (PDB ID 5M3T, chain B) state were performed using Chimera (UCSF) software with the structure editing tool [27]. We abolished the pH sensitivity of residues D139 and E188 by replacing them with N139 and Q188 without affecting their size and overall protein structure.

### D. Expression and purification of wild-type and mutant CD81_LEL_ constructs

The mutant hCD81_LEL_ genes were synthesised by GENSCRIPT (New Jersey, USA) and cloned into the pHLSec vector as previously done for the wild-type gene [18]. Cloning was done into the pHLSec vector using the AgeI and KpNI restriction sites. This digestion introduced in the protein sequence the extra residues ETG and GTKH6, respectively, at the N terminus and the C terminus. The corresponding three proteins were produced as secreted proteins by transient transfection of human embryonic kidney cells (HEK 293T) and purified as described elsewhere [18]. In brief, after IMAC purification using nickel-coated beads (CYTIVA, Massachusetts, USA), the protein was concentrated, buffer-exchanged into 20 mM Tris (pH 7.2) and 150 mM NaCl, and further purified by size-exclusion chromatography on a Superdex 75 10/300 Increase (CYTIVA, Massachusetts, USA) column. The eluted fractions corresponding to the wt-hCD81_LEL_ dimer were concentrated to ∼10 mg/mL using Vivaspin concentrator (MWCO 10,000; SARTORIUS, Göttingen, Germany) in 50 mM TRIS pH 7.4 and 200 mM NaCl, stored at 4°C and used within 48 h for CD experiments or flash-frozen in liquid nitrogen with the addition of 10% glycerol and stored at -80 ºC until further use. The mutant hCD81_LEL_ proteins were collected from the elution fractions and used fresh in a ∼1mg/mL concentration.

### E. Circular dichroism (CD) spectroscopy and protein stability analysis as a function of temperature

Prior CD experiments mutants and wild-type CD81_LEL_ samples were buffer exchanged using Micro Bio-Spin columns (BIORAD, California, USA) or Vivaspin (MWCO 10,000; SARTORIUS, Göttingen, Germany) into the following buffers: (i) 20 mM Sørensen phosphate pH 7.4, 50mM sodium sulphate; (ii) 20 mM phosphate-citrate pH 5.5, 50 mM sodium sulphate; (iii) 20 mM phosphate-citrate pH 4.6, 50 mM sodium sulphate. CD secondary structure spectra for all pH values were collected on a Jasco series J-815 spectropolarimeter before the thermal unfolding studies. Hellma-absorption cell (type 106-QS, light path length of 0.1 mm) (HELLMA, Buchs, Switzerland) was utilised to obtain the spectral data in the 190-250 nm regions for each buffer condition at 21 °C. Data points were collected every 0.2 nm at a scan rate of 50 nm/min. The concentration of the wild-type and mutant CD81_LEL_ proteins employed in these studies was estimated using a micro-volume Nanodrop spectrophotometer (0.7 mg/ml). The 6x-measurements averaged data with baseline adjustment were analysed using the neural network software K2d through the DICHROWEB server [28]. Then the thermal stability profile of each protein sample was determined by recording the intensity of ellipticity at 222 nm from 21 °C to 95 °C. The heating rate was 1o C/min with a scan rate of 60 °C/h. The temperature was incremented every 0.2 ºC. A Tectron-Bio 20 circulating water bath (JP SELECTA, Spain) was used as a thermostat for the optical cell’s Jasco CDF-426S Peltier temperature controller. These experiments were performed in duplicate/triplicate, and the average of the two experiments is shown in Table 1. The Tm of each construct at the different pHs was determined from fitting the curves of the thermal denaturation profiles to the two-state model described in the CD-Pal software [29].

**Table 1.** Biophysical data generated from CD-Pal analysis for wild-type and mutants CD81_LEL_ analysed by CD spectroscopy at different pHs.

| **CD81<sub>LEL</sub> pH 7.4** | ***n*** | **$T_M \pm e^*$** | **$\Delta H_{Tm} \pm e^*$** |
| --- | --- | --- | --- |
| wild-type | 2 | $79.07 \pm 0.25$ | $258.99 \pm 9.09$ |
| Mut1 D139N | 3 | $77.52 \pm 0.20$ | $295.52 \pm 9.18$ |
| Mut2 E188Q | 2 | $79.13 \pm 0.25$ | $308.56 \pm 8.52$ |
| **CD81<sub>LEL</sub> pH 5.5** |  |  |  |
| wild-type | 3 | $79.01 \pm 0.51$ | $252.36 \pm 12.44$ |
| Mut1 D139N | 2 | $77.85 \pm 0.16$ | $266.16 \pm 4.26$ |
| Mut2 E188Q | 2 | $78.89 \pm 0.30$ | $254.5 \pm 8.3$ |
| **CD81<sub>LEL</sub> pH 4.6** |  |  |  |
| wild-type | 3 | $70.88 \pm 0.74$ | $250.36 \pm 23.10$ |
| Mut1 D139N | 2 | $71.98 \pm 0.21$ | $311.93 \pm 17.21$ |
| Mut2 E188Q | 2 | $68.57 \pm 0.09$ | $225.73 \pm 2.98$ |

*n*: number of measured samples
*e*\*: jackknife error
*T<sub>m</sub>*: Average melting temperature
$\Delta H_{Tm}$ : Enthalpy of denaturation

## III. Results

### A. Changes in enthalpy of hydration drive the conformational change of CD81_LEL_

Previous simulations studies postulated that the head-subdomain CD81_LEL_ changes from the close to the open structure upon lowering the pH [18]. To understand the influence of pH on the structural plasticity of CD81_LEL_, we analysed how the pH affected the total net-charge of the protein. We explored at pHs: 7.4, 6.2, 5.5, 5.0 and 4.6; that mimic the pH conditions of the endosomal-lysosomal pathway [5,30].

We first performed an in-silico analysis identifying the ionisable groups/residues of the protein to investigate differences in the electrostatic potential of the CD81_LEL_ protein across the close and open conformations. Five residues, D139, H151, E152, E188, and H191 are pH-sensitive and influence the electrostatic potential of CD81_LEL_ at low pH (Fig. 1B). At low pH, the close conformer’s charge becomes neutral while the open conformer’s one becomes positive. Both have a negative net charge at physiological pH. Residues D139 and E188 are located on the protein’s surface, proximal to the head subdomain (helices B, C and D), while H151, H191 and E152 are located in the core of the large extracellular loop (Fig. 1B). These observations suggest that the changes in the electrostatic potential of CD81_LEL_ at these locations trigger the transition close-to-open CD81_LEL_ (Fig. 1C). To test this hypothesis and to assess what effect would have the titration of residues D139, H151, E152, E188 or H191 (Fig. 1C) on the protein’s stability (enthalpy); we monitored the changes occurred at a desired pH from the endosomal pathway, normalising the data to pH 7.4. In this way, the intra-protein’s (ΔH^P^_pH_= H^P^_pH_ − H^P^_pH 7.4_) and the protein-solvent’s (ΔH^PS^_pH_= H^PS^_pH_ − H^PS^_pH 7.4_) enthalpies were obtained. It is important to highlight how positive values of ΔH indicate an increase in the enthalpy and hence a destabilising effect. Conversely, negative values correspond to stabilising changes.

To this end, we performed a set of 2 ns molecular atomistic simulations in the close and open structures titrated at pHs 7.4, 6.2, 5.5, 5.0 and 4.6. The simulation time also permitted the sampling of the water configurations of the solvent. Changes in the enthalpy fully characterize the interactions between the molecule and the water [31–33]. As expected, we confirmed that the initial structure of the protein is preserved within 2 Å RMSD. The ΔH^P^ profiles in Fig. 2A of CD81_LEL_ close conformation do not show a destabilizing energy change as a function of pH. This result indicates that intra protein interactions are not responsible for the pH mediated conformational change (supplemental material Text-S1 and Fig. S1-S5).

**FIG 2.**
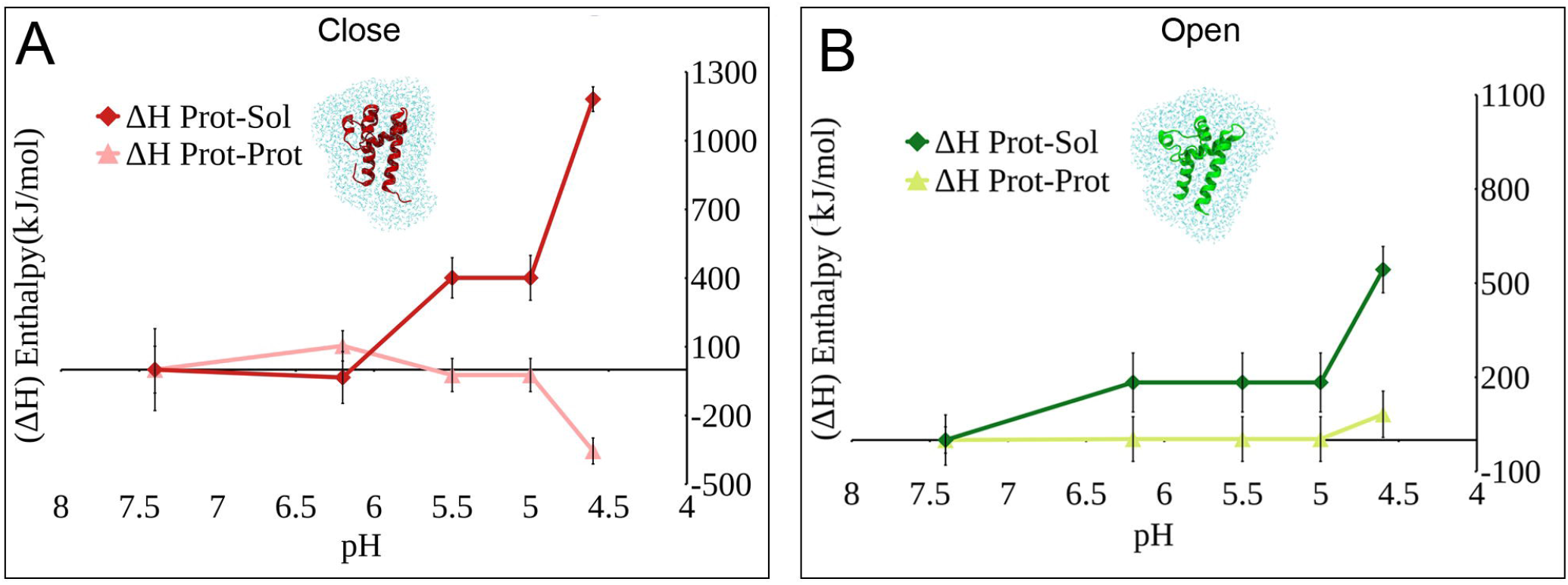
Change in enthalpy (ΔH) of CD81_LEL_ conformations at different endosomal pH. (A) Change in enthalpy (ΔH^PP^ and ΔH^PS^) of the CD81_LEL_ close. (B) Change in enthalpy (ΔH^PP^ and ΔH^PS^) of the CD81_LEL_ open. All energy values have been normalised against the reference state assumed at pH 7.4. The plots show that the protein-solvent interactions are the largest contribution to the destabilization of the closed conformer upon lowering pH.

In Fig. 2A, we show that there is a substantial increase in ΔH^PS^ upon lowering pH. Specifically, we found that the stability of the close structure is dominated by the hydration shell enthalpy, which stabilises the protein at pH 6.2 (Fig. 2A and Fig. S6). Below pH 6.2, the protein becomes unstable due to a destabilisation of the protein-solvent interactions (Fig. 2A and Fig. S6). For the open conformation, the hydration shell contribution to the protein stability is practically constant up to pH 5 (Fig. 2B and Fig. S6). Hence, when low pH is simulated, the stability of the protein in water is compromised because the protein-solvent interactions become unfavourable. From this, we speculate that the pH sensing mechanism is mediated by protein-solvent interactions. It is important to stress that the changes in enthalpy that we measured are averaged over the water configurations.

### B. The specific change in enthalpy of hydration of residues E188 and D139 evidence the conformational change at low pH

To identify which residues contributed the most to the destabilization of the closed conformer, we monitor the change in protein-solvent enthalpy ΔΔH= ΔHc^PSpH^-ΔHo^PSpH^ (Fig. 3A) between the open and close structures for each CD81_LEL_ residue and at the different endosomal pH. We found that titration of E188 at pH 5.5 has the most destabilising ΔΔH showing a very distinct peak when compared to the other residues. E188 is located in the loop connecting helices D and E (Fig. 1B) and exposed to the solvent. This E188 residue appears protonated in both the close and open conformers at pH 5.5. However, its interaction with the water is much more destabilizing in the close conformation at pH 5.5 than in the open conformation (ΔHc > ΔHo). At pH 4.6 it is residue D139 that shows a distinctive peak in the ΔΔH among the other residues (Fig. 3A). Whereas D139 titrates at low pH in CD81_LEL_ close, increasing its enthalpy in terms of protein-solvent interactions, it remains charged in CD81_LEL_ open, favouring its enthalpy of interaction with water (Fig. 3B). D139 located in the loop connecting helices B and A, totally exposed to the solvent (Fig. 1B). The individual variation in enthalpy (ΔΔH) of interaction between residues H191, H151 or E152 with the solvent did not display significant differences across conformations (Fig. 3A), suggesting that the increase of enthalpy on CD81_LEL_ conformers in water is mainly driven by the titration of residues E188 and D139.

**FIG 3.**
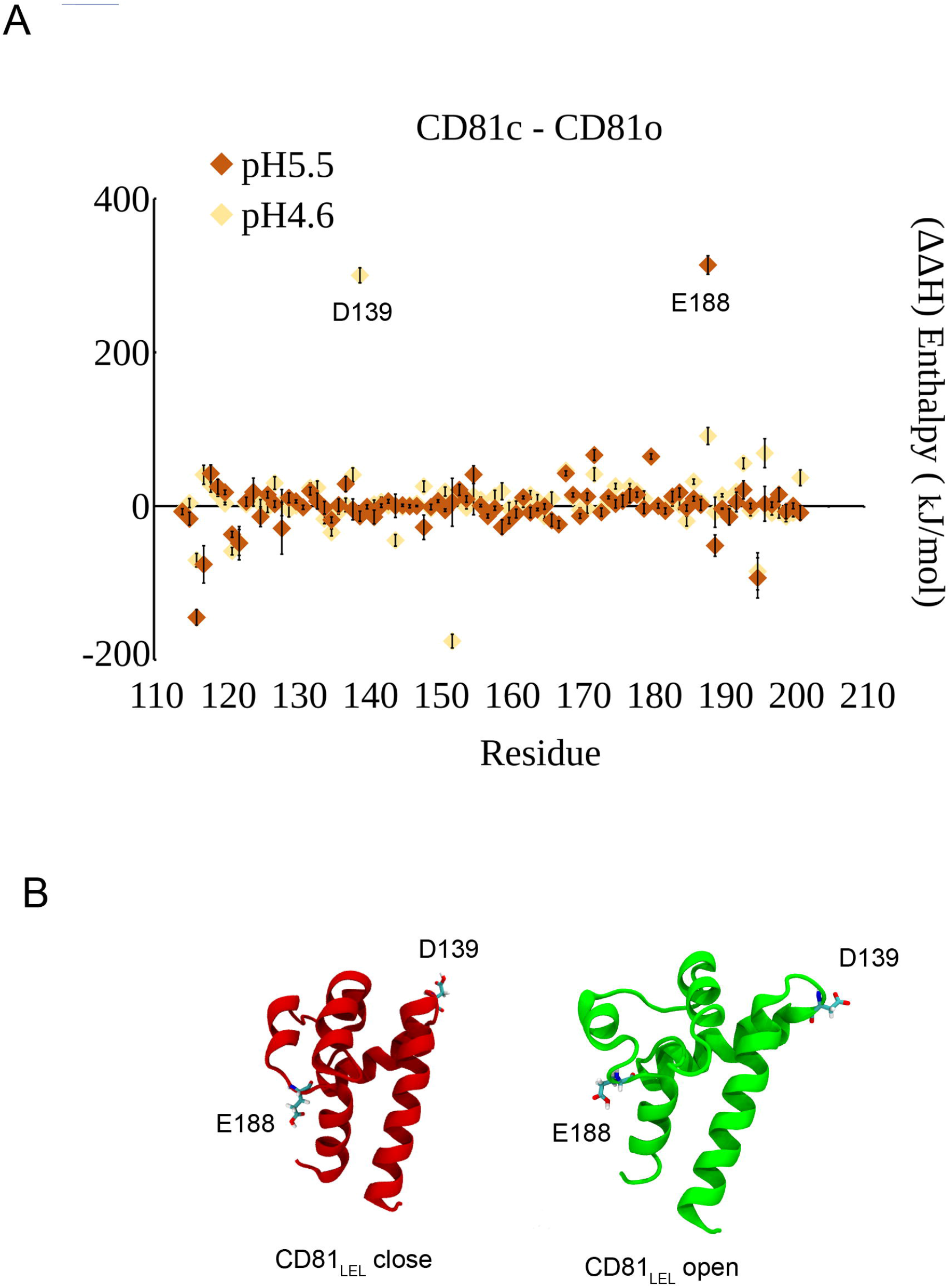
Protein-Solvent interactions by residue between CD81_LEL_ close and open conformations. (A) Differential enthalpy ΔΔH^PS^ of the protein-solvent interactions of each residue (112-204) of CD81_LEL_ at pH5.5 and pH 4.6. At pH 5.5 the residue E188 stands out with an isolated positive peak indicating a clear and isolated contribution to the destabilization. Lowering further the pH at 4.6 the peak of E188 is replaced by a new one of D139 equally distinctive. (B) Location and titration state of residues D139 and E188 on CD81_LEL_ close and open (PDB IDs 5M3T and 5M33).

### C. The solvation profile of residues E188 and D139 at low pH influences the stability of the different conformers of CD81_LEL_ at low pH

To elucidate the relationship between the protein-solvent interactions and the pH, we first inspected the distribution of solvent molecules around residues D139 and E188, responsible for the energetic changes. We used the Radial Pair Distribution Function, g(r), to quantify the number of water molecules around an amino acid. The g(r) is defined as the number of molecules found in the spherical shell at a distance r from the reference amino acid, normalised by the volume of the spherical shell.

Analysis of the average distribution of water around the residues D139 or E188 by the g(r) function showed a significant variation in the solvation profile of positions E188 and D139 on the CD81_LEL_ open-close structures upon varying the pH values (7.4-4.6) (Fig. 4 and Fig. S7). In the close conformer, D139 decreases the hydration level when simulating acidic conditions compared to the open conformation, in which the water layer remains constant. For E188, both open and close conformations experience a decrease in the hydration level at this position at low pH. These results are corroborated by a hydrogen bond analysis which shows a similar decrease in the hydrogen bond profile of residues E188 or D139 when comparing the close conformer of CD81_LEL_ close and open at physiological and low pH (Fig. S8).

**FIG 4.**
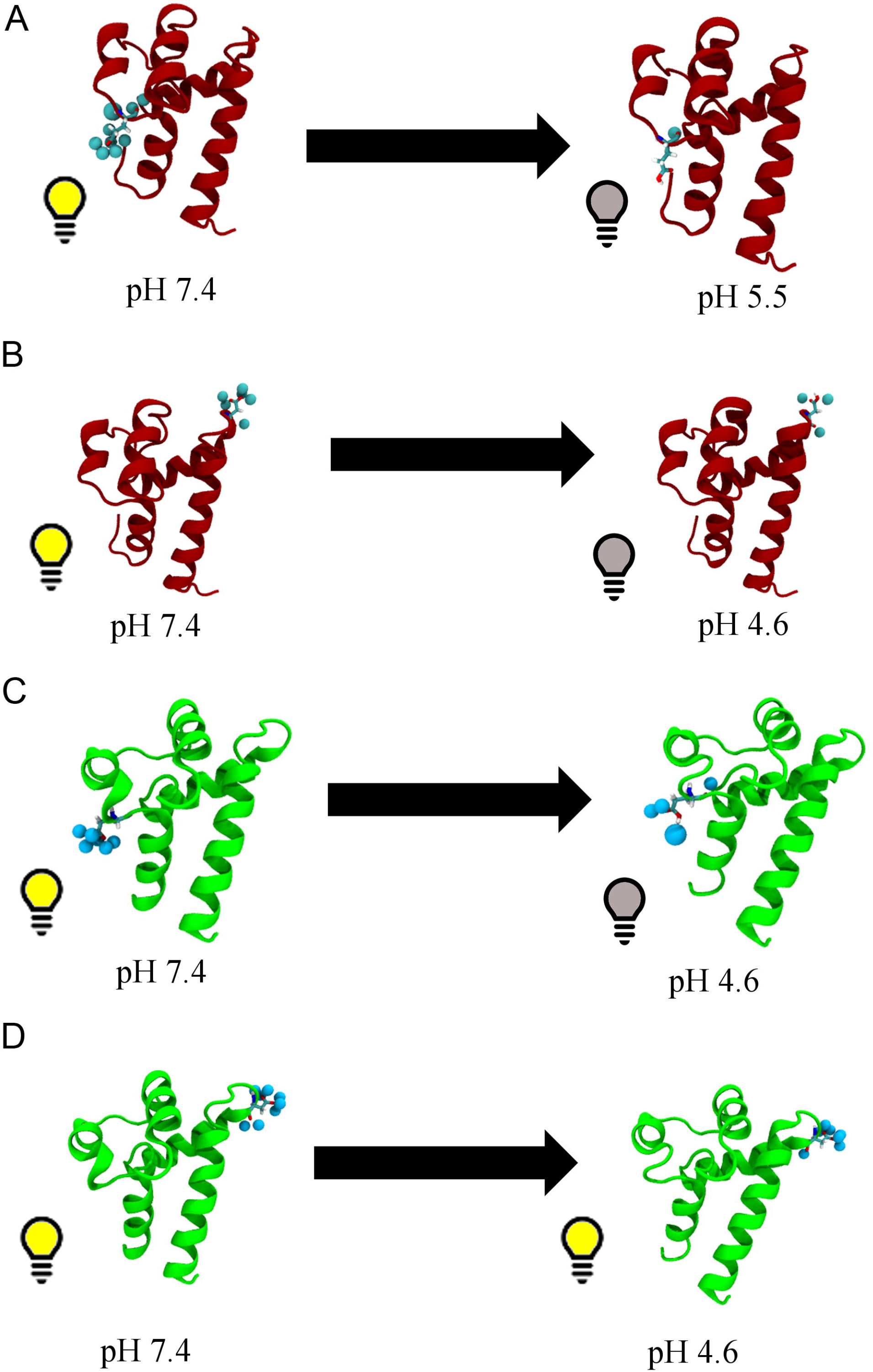
The stability of the CD81_LEL_ conformers is dictated by the titratable state of residues D139 and E188. (A-B) E188 and D139 titrate (switch off their charge) at low pH 5.5 and 4.6 respectively, losing interactions with water. This causes the close CD81_LEL_ conformation lose stability in water (ΔH>0) at acidic conditions; (C) In CD81_LEL_ open, E188 delays the switching off of its charge to lower pH 4.6 than in the close conformation, pH 5.5; stabilizing the open conformation at low pH. (D) D139 maintains its charge on at low pH in CD81_LEL_ open, making that conformation stable at acid pH conditions. Water molecules at 2 Å distance are depicted as cyan spheres.

Our results indicate that the titratable state of residues E188 and D139 regulate the amount of solvent around the head subdomain and, consequently, influence the stability of the different conformers of CD81_LEL_ at low pH.

### D. Mutations E188Q and D139N stabilise the close conformation of CD81_LEL_ in water by retaining the solvation profile at low pH

To test the involvement of residues E188 and D139 in regulating the CD81_LEL_ solvation, we mutated the E188 and D139 to not pH-sensitive residues. We chose the mutations E188Q and D139N that maintain the size and polar interactions of CD81_LEL_ with water but remove their pH dependence.

Following our enthalpy calculation protocol, we analysed the ΔH of the intra-protein enthalpy (H^P^) and the protein-solvent enthalpy (H^PS^) after mutating residues E188 or D139 in CD81_LEL_ close and CD81_LEL_ open. We found that mutations E188Q or D139N restored the stability of the CD81_LEL_ close conformation within the water at low pH (ΔH^PS^), compared to the wild-type (wt) system (Fig. 5A). Moreover, mutations E188Q and D139N have little influence on the enthalpy of the open conformation of CD81_LEL_ within the solvent (Fig. 5B).

**FIG 5.**
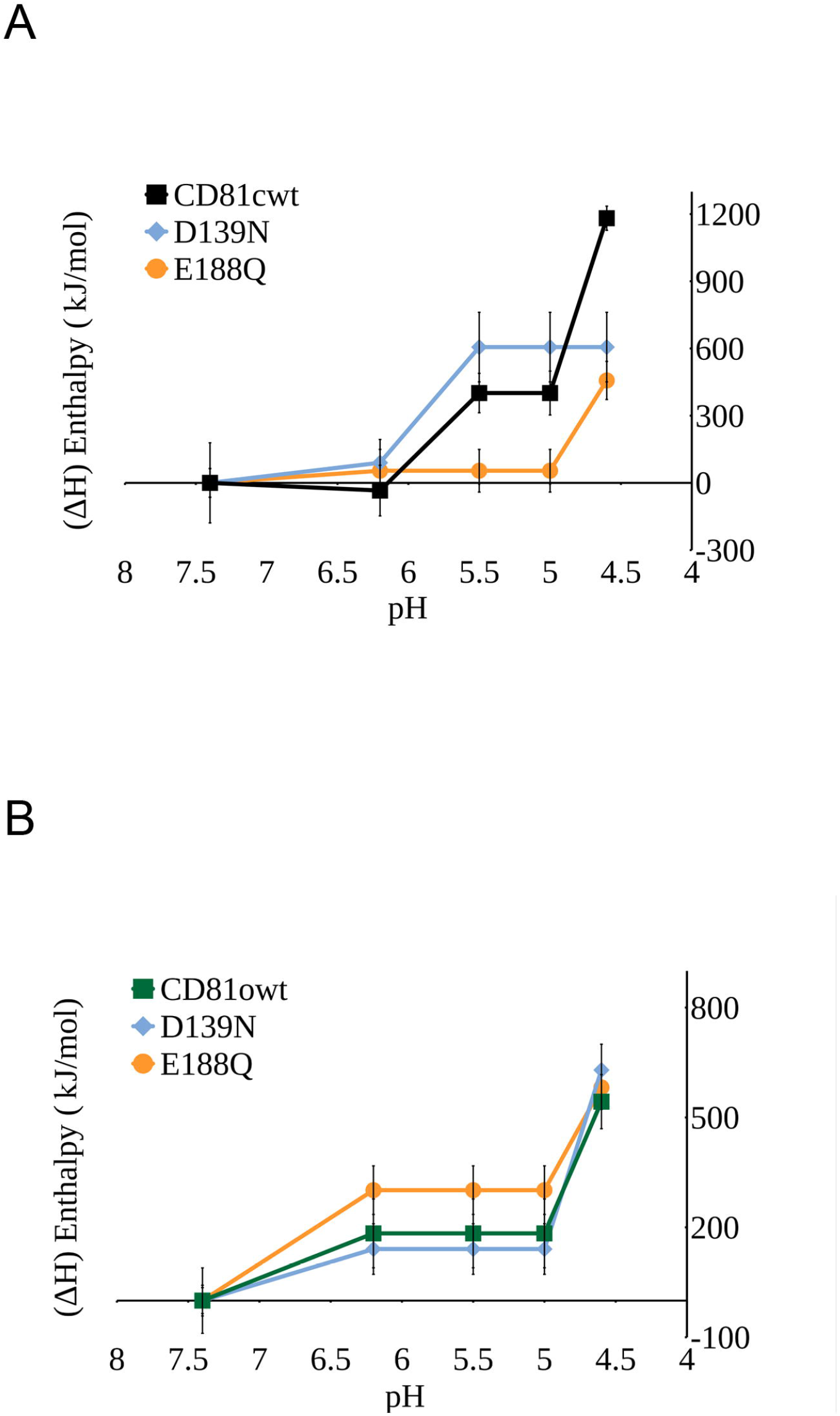
Evolution of the enthalpy of the protein-solvent (ΔH^PS^) interactions of CD81_LEL_ mutated at different endosomal pH. (A) At acidic conditions (pH4.6), E188Q or D139N stabilise the close structure’s enthalpy within water compared to the wild-type system. (B) Mutations E188Q or D139N have an imperceptible effect over the open structure’s enthalpy within water compared to the wild-type system.

In parallel, analysis of the average water distribution among residues N139 or Q188 by the g(r) function showed that mutations E188Q and D139N do not vary the system’s solvation profile during endosomal titration. These data support our hypothesis that the ionisable state of positions E188 and D139 influences the stability, in terms of enthalpy, of CD81_LEL_ conformers at low pH by modulating the liquid shell surrounding the molecule (Fig. S9).

### E. In silico mutations of E188Q or D139N shift the equilibrium towards the close conformation of CD81_LEL_ but barely affect the dynamics of CD81_LEL_ open

To study the effect of residues E188 and D139 on the flexibility of CD81_LEL_, we performed 1 µs molecular dynamics experiments on CD81_LEL_ close and open at pH 5.5 or pH 4.6, pH values at which positions E188 or D139 titrate, respectively.

At pH 5.5 we observe that minor oscillations in the dynamics of the head subdomain of wild-type CD81_LEL_ in close conformation, with an overall RMSD of 2-3 Å from the reference crystal structure (PDB ID 5M3T, chain B) (Fig. 6A and Fig. S10A). Mutations E188Q and D139N stabilize the dynamics to the observed crystallographic conformation (PDB ID 5M3T) of the protein by shifting the equilibrium configuration to RMSD values around 1 Å (Fig. 7A and Fig. S10A).

**FIG 6.**
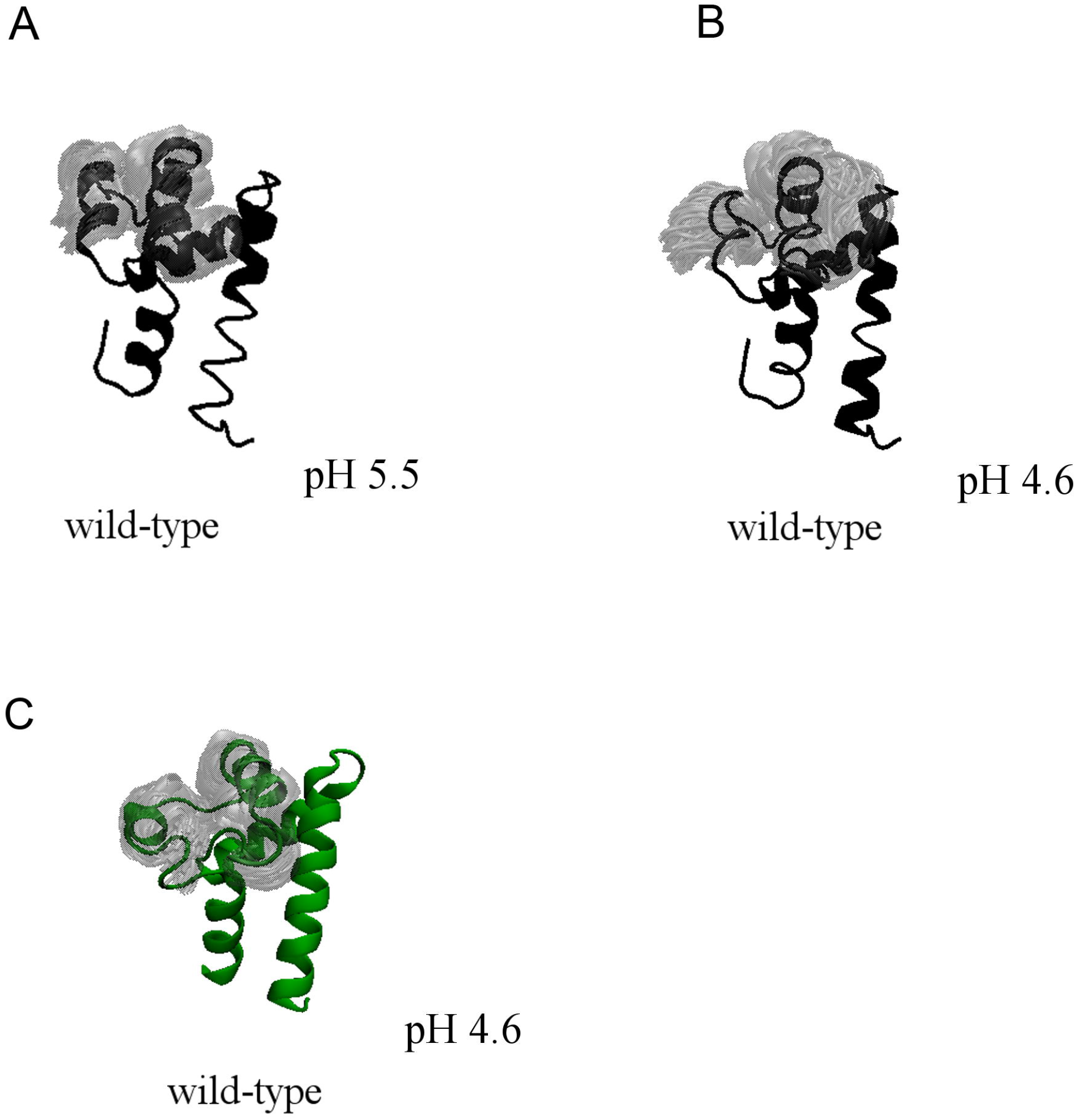
Dynamics of the head subdomain of CD81_LEL_. (A) Dynamics of the wild-type CD81_LEL_ at pH 5.5 and pH 4.6 in close conformation; (B) Dynamics of the wild-type CD81_LEL_ at pH 4.6 in open conformation. The grey shade indicates the extension of the fluctuations. Larger the shade larger the fluctuations. We quantified the fluctuations in Figure S10.

**FIG 7.**
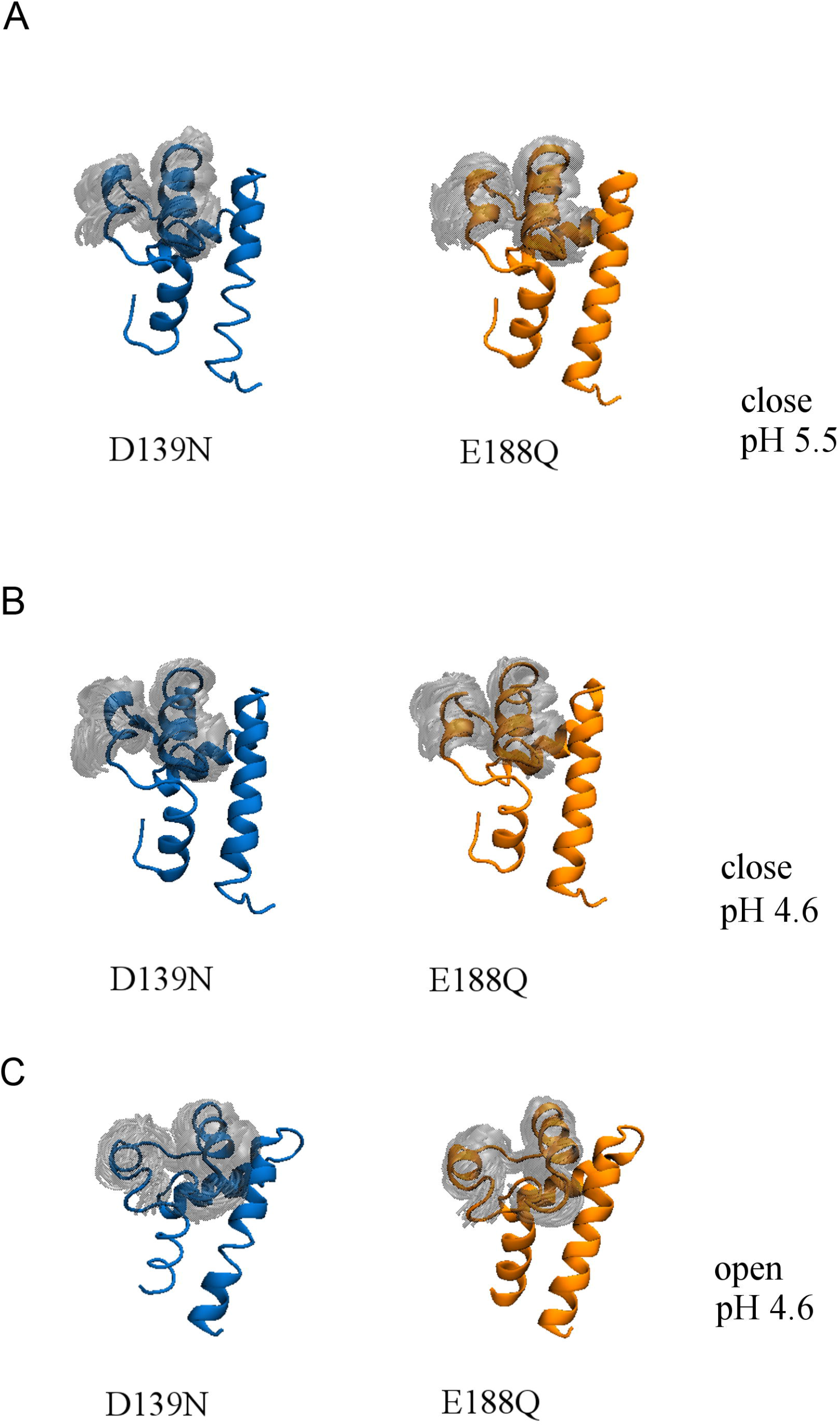
Dynamics of the mutant constructs of the head subdomain of CD81_LEL_. (A) Dynamics of the mutant constructs (D139 or E188) at pH 5.5 and (B) pH 4.6 in the close conformation. (C) Dynamics of the mutant constructs (D139 or E188) at pH 4.6 in the open conformation. The grey shade indicates the extension of the fluctuations. Larger the shade larger the fluctuations. We quantified the fluctuations in Figure S10 and S11.

When we simulated acidic conditions (pH 4.6), we observe the concomitant opening events on the wild-type form of CD81_LEL_, with a displacement of the equilibrium configuration towards RMSD values above 4 Å (Fig. 6B and Fig. S10B) from the reference crystallographic observed conformation. This shift suggests that titration of both positions E188 and D139 is necessary to free the movement of the head subdomain. Mutations E188Q and D139N stabilise the head subdomain’s compact form, as we observe a shift in the equilibrium configuration to RMSD values around 1 Å (Fig. 7B and Fig. S10B).

Our 1 µs molecular dynamics experiments on the open conformer of CD81_LEL_ correlate with previous studies carried out by Cunha *et al*. 2017 [18] - the open state of the head subdomain of CD81_LEL_ is maintained in acidic conditions along 1 μs MD trajectory. The head subdomain of the protein experiences a minor fluctuation in its dynamic oscillations (Fig. 6C and Fig. S11A). Mutations E188Q or D139N have little effect on the plasticity of the head subdomain of CD81_LEL_ open as the dynamics explored by the wild-type and the mutated systems are equivalent (Fig. 7C and Fig. S11A).

However, we also observe a small population that drifts to RMSD values of 4 Å when we simulate CD81_LEL_ open with mutation E188Q (Fig. 7C and Fig. S11B). Comparing with the reference close conformation of CD81_LEL_ (PDB ID 5M3T, chain B) we observe that E188Q mutation is not enough to shift the equilibrium of the open conformation to the close structure (Fig. 7C and Fig. S11B).

### F. Residues E188 and D139 induce a global redistribution of water molecules around the protein that evidence the stability of the different conformations of CD81_LEL_

To further understand the role of residues E188 and D139 in the plasticity of CD81_LEL_, we assessed the effect of the in-silico mutations in the rest of the residues of the head-subdomain. To this end, we calculated an ΔΔH comparing the corresponding ΔH^PS^ of each residue of the mutated systems E188Q or D139N to the wild-type system at low pH, i.e.: ΔΔH^PS^ = ΔH^PS E188Q pH4.6^ – ΔH^PS wt pH4.6^. In Fig. 8 we plot individual ΔΔHs as a function of distance from D139 and E188 at pH 4.6.

**FIG 8.**
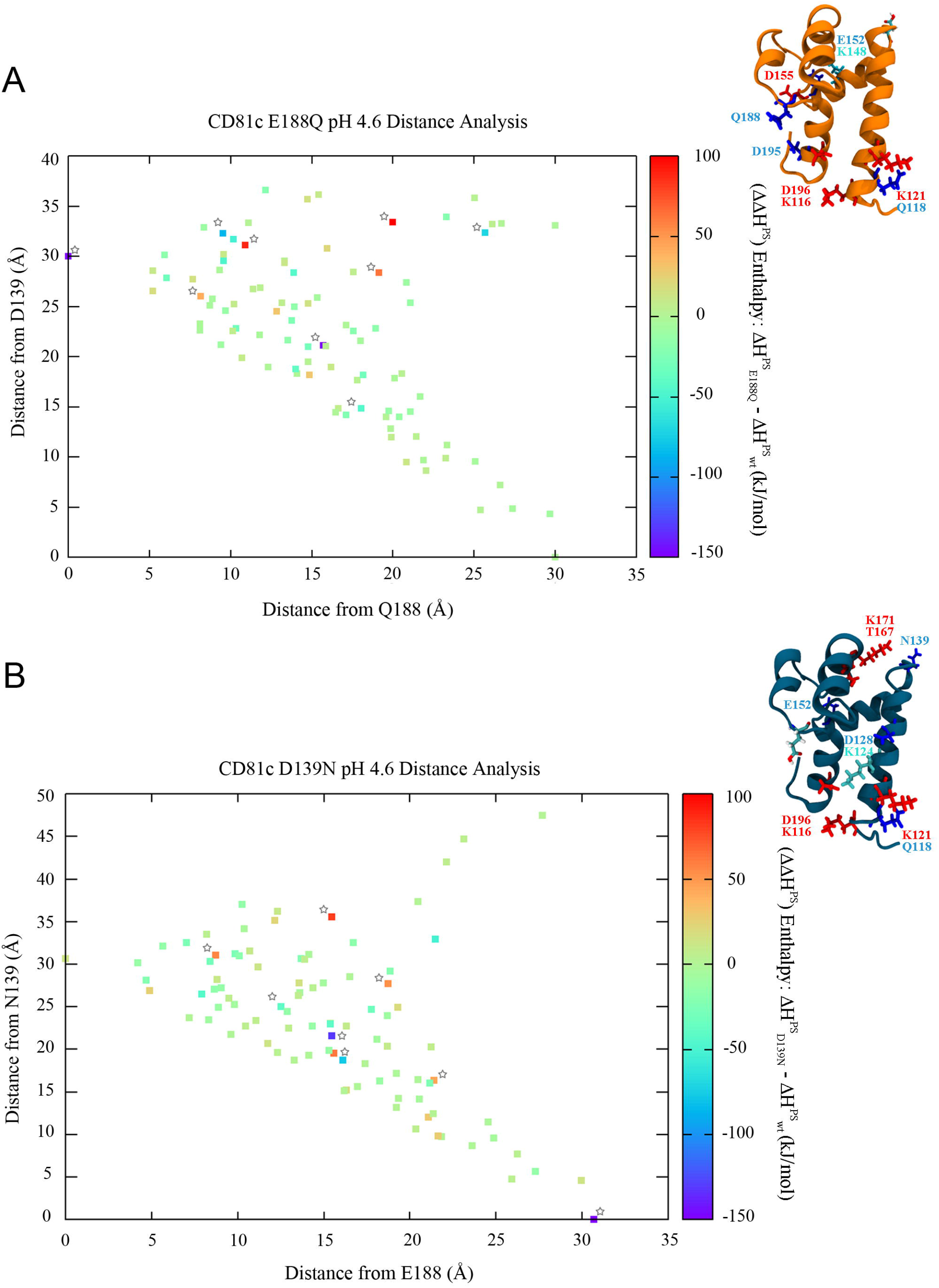
Distance Analysis on CD81_LEL_ close. (A) ΔΔH^PS^ of CD81^LEL^ close with mutation E188Q. (B) ΔΔH^PS^ of CD81_LEL_ close with mutation D139N. Stars highlight the position of the residues that get stabilised (blue) and destabilised (red), which are depicted as sticks in the cartoon protein structures.

The first distinguishable result is that the effect of the E188Q or D139N mutations is energetically neutral in the complete set of residues that form the head subdomain (Fig. 8). This further supports our initial observation that the conformational change is driven exclusively by the change in the solvation of the E188 and D139 residues at pH 5.5 and 4.6, respectively. A second outcome from this analysis is that the mutations affect the protein-solvent interactions of residues within a range of 5-20 Å distance from E188Q and within a range of 15-35 Å distance from D139N for both systems CD81_LEL_ close (Fig. 8) and CD81_LEL_ open (supplemental material Text-S1). In particular, in the CD81_LEL_ close conformation we observe that the mutation E188Q has a robust destabilising effect of the protein-solvent interactions (ΔΔH>50 kJ/mol) of residues K116 (20 Å), K121 (19 Å), D155 (8 Å) and D195 (10 Å). In contrast, Q118 (25.7 Å), E152 (15.7 Å), D196 (9.5 Å) get their interaction with water stabilised (ΔΔH<-50 kJ/mol) (Fig. 8A).

In the case of mutation D139N, the profile of residues affected is similar: K116 (35.6 Å), K121 (27.7 Å), T167 (19.5 Å), T171 (16.3 Å) and D195 (31 Å) whose interactions with water become unstable (ΔΔH> 50 kJ/mol) while the interaction of residues Q118 (32.9 Å), D128 (18.7 Å), and E152 (21.6 Å) with water become stable at low pH (ΔΔH<-50 kJ/mol) (Fig. 8B).

### G. Role of the solvent combined enthalpic and entropic contributions

So far, we explored the role of the enthalpy inducing the conformational change and as shown previously, both open and closed structures have similar dynamic oscillations (see Fig. S12). Therefore, any significant entropic contribution should come from the solvation entropy of the water. To this end, we measured the probability of finding a water molecule close to the residues, a quantity that is equivalent to the solvation free energy. The probability of finding a water molecule at a distance r from a residue has been estimated through the radial distribution function g(r) described in the method sections. The g(r) analysis confirms that positive changes ΔΔH>0 in enthalpy correspond, in general, to a loss of water solvation. Conversely, negative changes ΔΔH<0 in enthalpy correspond to a gain of water solvation (supplemental material Text S1 and Fig. S2-S3). It is important to highlight that the g(r) is the probability density of finding a water molecule at a distance *r* from a specific residue. By considering the Boltzmann definition of the free energy F(r) of a water molecule at distance r

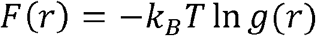

we see the direct dependence of the free energy from the g(r). Hence, the g(r) analysis takes into account both the enthalpic and entropic contribution coming from the solvation. In particular, the analysis over the solvation g(r) indicates that the signal triggered by residues D139 and E188 to regulate CD81_LEL_ plasticity propagates around the whole head subdomain at low pH.

### H. Protein stability of wild-type and mutants CD81_LEL_ by circular dichroism

To assess experimentally the impact of the suggested mutations by the in silico mutational study we expressed and purified the wild-type and mutant proteins in the HEK293T mammalian cell system (Fig. S13) and performed circular dichroism spectroscopy analysis. Size-exclusion chromatography during protein purification (Fig. S13) indicated that the mutant proteins form dimers as the wild-type, supporting the fact that the introduced mutations do not alter the ability of dimerization. Three different buffers at pH 7.4, pH 5.5 and pH 4.6 were used to assess the thermal stability of the wild-type and mutant proteins. At the given pHs, overlaid CD Spectra (190–250 nm) of CD81_LEL_ wild-type, D139N (mut-1), and E188Q (mut-2) indicated that each protein was fully folded and displayed α-helix secondary structure (Fig. S14A). Prediction of the secondary structure CD spectra of both open and close conformations in monomeric or dimeric form using PDB2CD software showed marginal differences among the spectra [34] (Fig. S14B).

Thermal stability analyses were performed at a fixed wavelength (222 nm) at a temperature interval of 21 ºC – 95 ºC at the different pHs mentioned: pH 7.4, pH 5.5 and pH 4.6. At pH 7.4, the three proteins, wild-type, and mutants, showed a transition from folded to unfolded at temperatures above 75 ºC (Table 1). We observe a decrease in the ellipticity measured above this temperature (Fig. S15). At pH 5.5, the three proteins display a similar behaviour. The transition from folded to unfolded occurs at temperatures above 75 ºC, suggesting that moderate acidic pH does not affect the overall thermal stability of the wild-type and mutant CD81_LEL_ recombinant proteins (Table 1; Fig. S16). However, at low pH 4.6, the overall thermal stability of the wild-type and mutant proteins is compromised. The CD spectra show a decrease in the melting temperature up to 10 ºC (Table 1) for the transition from folded to unfolded in the three constructs (Table 1; Fig. S17).

Nevertheless, if we compare the extrapolated enthalpy of denaturation (ΔH_Tm_) at the studied pHs, we note subtle differences between the three systems, wild-type, mut-1 (D139N) and mut-2 (E188Q). At physiological pH, the mutations stabilise the protein, as more energy is required for denaturalisation (Table 1). At pH 5.5, on the contrary, the three proteins behave similarly, suggesting that moderate acidic pH does not significantly affect the overall thermal stability of the system. At low acidic pH 4.6, the denaturation enthalpy is significantly different among the three proteins studied, being CD81_LEL_ carrying mutation D139N the most stable system.

## IV. Discussion

Our full atomistic molecular dynamics simulations have shown that the conformational plasticity of the head subdomain of CD81_LEL_ is regulated by positions D139 and E188. By doing a comparative analysis with the mutated (D139N and E188Q) systems, we have been able to identify that the titration of these residues in the close conformation of CD81_LEL_ at low pH disrupts the head subdomain’s solvation profile. Charge neutralisation at positions E188 and D139 due to their titration at low pH leads to a relaxation of the H-bond network at the protein-solvent interface, facilitating the head subdomain dynamics to adopt the open conformation, the latter more stable at low pH than the close conformation (Fig. S8).

Secondary structure analysis on the different constructs expressed (wild-type, mut-1, mut-2) by CD spectra scanning have revealed no significant differences in α-helical content between wild-type or mutated proteins, indicating that the mutants are fully folded. From the CD analysis we compared the thermal stability of wild-type proteins with the mutants. The experiments shown that the mutation D139N increases the protein’s stability at low pH, as the enthalpy for denaturation of the system is greater than that of the wild-type and mut-2 systems (Table 1). This result would confirm that titration of residue D139 governs the dynamics of the solvent around CD81_LEL_ at low pH. As illustrated in Figure 8 the distance analysis points out that more residues are affected by mutation D139N than E188Q. This supports our initial hypothesis that the conformational transition from close to open in CD81_LEL_ is triggered by the titration of E188 that then is strongly amplified by the effect of titration of D139.

Considering previous studies [18] and our findings, the molecular mechanisms involved in the regulation of the conformational plasticity of CD81_LEL_ could be driven by a combined signal. First, by the sequential titration of residues E188 and D139 at pH 5.5 and pH 4.6, and secondly by the titration of residue H151, in the core of the head subdomain, which amplifies the effect of the distal residues. At low pH, the disruption of the disulphide bond C157-C175 may also contribute to the system’s opening, as shown previously [18]. However, in our MD studies, the two disulphide bonds (C160-C190; C157-C175) are preserved, suggesting that the presence of the disulphide bridges is not a structural factor determinant for the plasticity of CD81_LEL_.

Moreover, variation across species of the residue E188 correlates to the organism susceptibility or not to HCV infection. This susceptibility is determined by the ability in some organisms of E188 to lock CD81_LEL_’s closed structure by forming hydrogen bonds or salt bridges with the residue at position 196. This is a structural feature unavailable in the human CD81_LEL_ [35]. This premise suggests the role of residue E188 in regulating the CD81_LEL_ plasticity.

Overall, our study shows that the solvent microenvironment surrounding CD81_LEL_ is the crucial player in the plasticity of the head-subdomain. Residues E188 and D139 act as pH sensors in the CD81_LEL_ close conformation by triggering the change towards the open conformation by modulating the distribution of the surrounding water molecules in the head subdomain. Such a mechanism is distinct from the ones presented in previous studies focusing on the role of short and long-distance water structure on the dynamics of folding, enzymatic activity, or protein plasticity. Earlier findings by Dahanayake *et al*. 2018 [36] and Gavrilov *et al*. 2017 [37] demonstrate how the protein dynamics depend on the local solvation environment’s friction, also named solvent viscosity. This property is directly related to the solvent shell dynamics in which the protein’s motion is coupled with the translational motion of the surrounding shell water. Water friction can be modulated with pH or ionic strength. The chemical nature of the exposed amino acids is modified, and therefore the solvation entropy and solvation free energy is altered [37]. This dynamic effect results in a reorientation of the water molecules of the solvation shell around the protein, allowing conformational changes in the protein’s structure itself, *i*.*e*. proton channel opening triggered by the protonation of histidine tetrads. This titration disrupts the H-bond network between these histidine residues, allowing proton transport at low pH in Influenza A viruses to mediate fusion [38–41]. Hence, our equilibrium mechanism for the CD81_LEL_ is an additional piece of the allosteric transition puzzle complementing the known dynamical scenario [42]. The mechanism proposed here is a different form of solvent-induced allosteric transition in proteins, in which the signal triggered by distal residues D139 and E188 is translocated through the surrounding solvent to the rest of the protein, allowing the opening of the head subdomain. We propose that this allosteric mechanism could serve as a pH sensing strategy for CD81 to time biological signals along endosomal pathway [11]. At infection, HCV bound to the CD81 receptor could exploit this molecular reactivity to sense the virus-cell entry steps, from viral attachment at physiological pH to the maturation state of the endosomes with the pH drop. The CD81_LEL_ conformational transition from pH 5.5 to pH 4.6 could drive the structural changes of the viral glycoproteins as reported by [12] and set the time for fusion in the endosomes.

## Acknowledgements

We are grateful to Ane Martinez-Castillo (Abrescia Lab) and Nicanor Zalba Olaizola (Millet Lab) for valuable discussions on protein expression and purification, and on circular dichroism, respectively. We are also in debt to Francisco J. Blanco and Tammo Diercks for a preliminary discussion on spectroscopic protocols.

## Disclosure statement

No potential conflict of interest reported by the authors

## Funding

We acknowledge the computer resources at Tirant and technical support provided by the Spanish Supercomputing Network (BCV-2019-2-0007). We also thank the support of the computing infrastructure of the i2BASQUE academic network. This research was supported by the Spanish Ministerio de Ciencia, Innovación y Universidades (RTI2018-095700-B-I00 to N.G.A.A.), by the CIC bioGUNE and CIC biomaGUNE (CICBMG-PhD-2018-02296 to C.R.F.). This research was supported by the Spanish Ministerio de Economía y Competitividad (MINECO) (FIS2017-89471-R to I.C.). This research was supported by Programa Red Guipuzcoana de Ciencia, Tecnología e Información (2019-CIEN-000051-01 to I.C.). I.C. acknowledges support from try BIKAINTEK program (grant No. 008-B1/2020). MICINN is also thanked for the Severo Ochoa Excellence Accreditation to the CIC bioGUNE (SEV-2016-0644) and the María de Maeztu Units of Excellence Programme – Grant No. MDM-2017-0720 Ministry of Science, Innovation and Universities.

## Notes

### Competing Interest Statement

The authors have declared no competing interest.

