## Supplementary material for "Solvent-induced allosteric transition of the Hepatitis C virus human cellular receptor CD81 large extracellular loop": Supplmental Data

Supplemental Text-S1

Supplemental Fig. S1-S13

Supplemental References

**Supplemental Text-S1**

**Results**

***Relation between the ionizable profiles and the plasticity of CD81$_{LEL}$ at low pH***

To pinpoint which residues among the D139, H151, E152, E188 or H191 were relevant to make one conformation preferable versus the other we swapped the titration profiles for the close and the open conformations at the individual pHs (Fig. S1A). We observed an increase in the differential enthalpy ($\Delta H^{PS}$) of the interaction of the CD81$_{LEL}$ conformers with swapped net-charges in the water molecules at low pH (pH4.6) similar to that when the protonation states are not swapped (Fig. S1B). Through this analysis we noticed that among all the conformations, residues E188 and H191 appeared neutral at low pH, suggesting that titration of these specific positions, E188 and H191, could be playing a role in the stability of CD81$_{LEL}$ in water.

***Local changes in the water distribution around CD81$_{LEL}$ driven by E188Q and D139N mutations stabilise the close conformation at low pH***

Radial pair distribution function g(r) analysis revealed a redistribution of the average water molecules around the protein between the first (2 Å) and second solvation layer (3 Å) when the E188Q mutation was introduced. While some residues, K116, D155 and D195 lose water molecules others such as Q118, K121, E152, Q188 and D195 become more hydrated. In particular residue Q188 retains the solvent viscosity at this position, and thus, stabilize the close conformation of the CD81$_{LEL}$ (Fig. S2). In the case of mutation D139N, the analysis of the radial pair distribution function g(r) revealed a similar behavior of water molecules around the protein: a gain of water molecules is observed at residues Q118, E152 and K171 and a loss in the first and second solvation layer is noticed around residues K116, K121, D128, T167 and D195, as seen for the residues affected by the E188Q mutation (Fig. S2 and S3A). Moreover, although the level of hydration at N139 at low pH does not increase with respect to the wild-type system (Fig. S3B), we removed the pH dependence of this residue, blocking changes on its hydration shell as effect of pH.

***Local changes in the water distribution around $CD81_{LEL}$ driven by E188Q and D139N mutations destabilise the open conformation at low pH.***

For $CD81_{LEL}$ open, the distance analysis revealed different effects of mutations E188Q and D139N on the protein. Although the effect of both substitutions is energetically neutral in the complete set of residues that form the head subdomain, as observed in $CD81_{LEL}$ close; the scope of E188Q mutation reaches a greater set of residues, as we observe a higher number of $\Delta\Delta H>0$ spots in the distance analysis graph (Fig. S4). In this particular mutation, E188Q, the strongest effects ($\Delta\Delta H>50$ kJ/mol) occur on residues D128, N133 and D196 at low pH; and the residues that have the greatest gain in enthalpy ($\Delta\Delta H<-50$ kJ/mol) are K144, K148 and Q188 (Figure S6A). Analysis of the radial pair distribution function g(r) revealed a loss of hydration on those residues negatively affected by the E188Q mutation (D128, N133 and D196) (Fig. S5A). These water molecules redistribute around the protein, stabilizing in particular, residues K144, K148 and Q188 (Fig. S5B). Q188, as observed in the close conformation, loses its pH dependence, stabilizing the open structure of $CD81_{LEL}$. In the case of mutation D139N, few residues are affected (Fig. S4). In particular, only residue D128 becomes unstable in water at low pH ($\Delta\Delta H>50$ kJ/mol) by losing water molecules on its solvation shell while E152 gets stabilized by acquiring a higher number of water molecules in its second solvation layer (Fig. S5A-B).

[3] T. Williams, C. Kelley, H.-B. Bröker, J. Campbell, R. Cunningham, D. Denholm, G. Elber, R. Fearick, C. Grammes, L. Hart, L. Hecking, P. Juhász, T. Koenig, D. Kotz, E. Kubaitis, R. Lang, T. Lecomte, A. Lehmann, A. Mai, B. Märkisch, E. A. Merritt, P. Mikulík, C. Steger, S. Takeno, T. Tkacik, J. Van Der Woude, J. R. Van Zandt, A. Woo, and J. Zellner, *Gnuplot 4.6 An Interactive Plotting Program*, (n.d.).

[4] E. S. Cunha, P. Sfriso, A. L. Rojas, A. Hospital, M. Orozco, and N. G. A. Abrescia, *Mechanism of Structural Tuning of the Hepatitis C Virus Human Cellular Receptor CD81 Large Extracellular Loop*, Structure **25**, 53 (2017).

[5] M. Niklasson, C. Andresen, S. Helander, M. G. L. Roth, A. Zimdahl Kahlin, M. Lindqvist Appell, and P. Lundström, *Robust and Convenient Analysis of Protein Thermal and Chemical Stability*, Protein Sci. **24**, 2055 (2015).

**FIGURES**

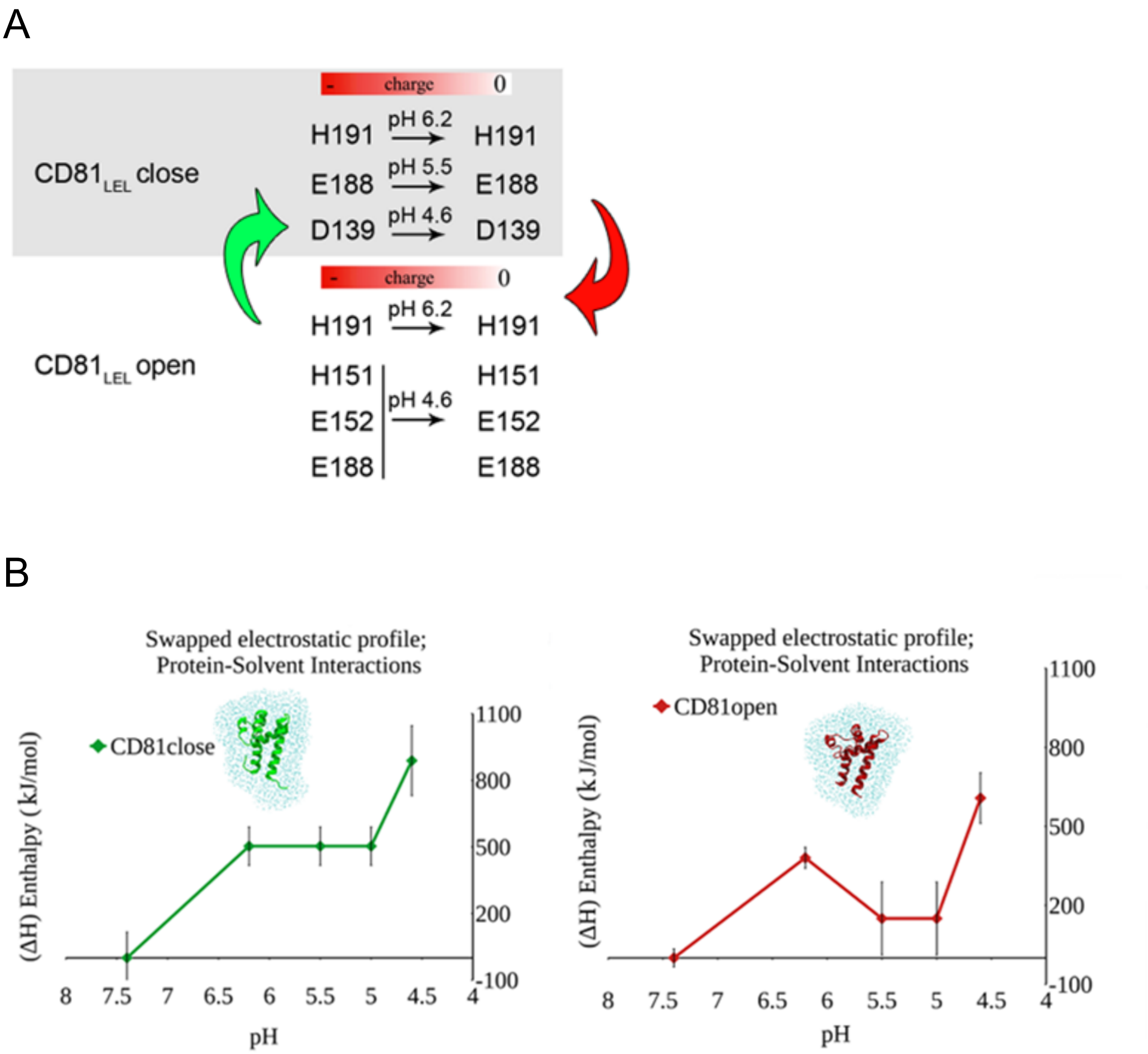


**FIG S1.** (A) Artificial profile of the $CD81_{LEL}$ conformers at low pH. Common pH dependent residues between conformations are highlighted in bold. (B) Difference of $\Delta H^{PS}$ of $CD81_{LEL}$ close and open but interchanging their pH dependent residues.

.

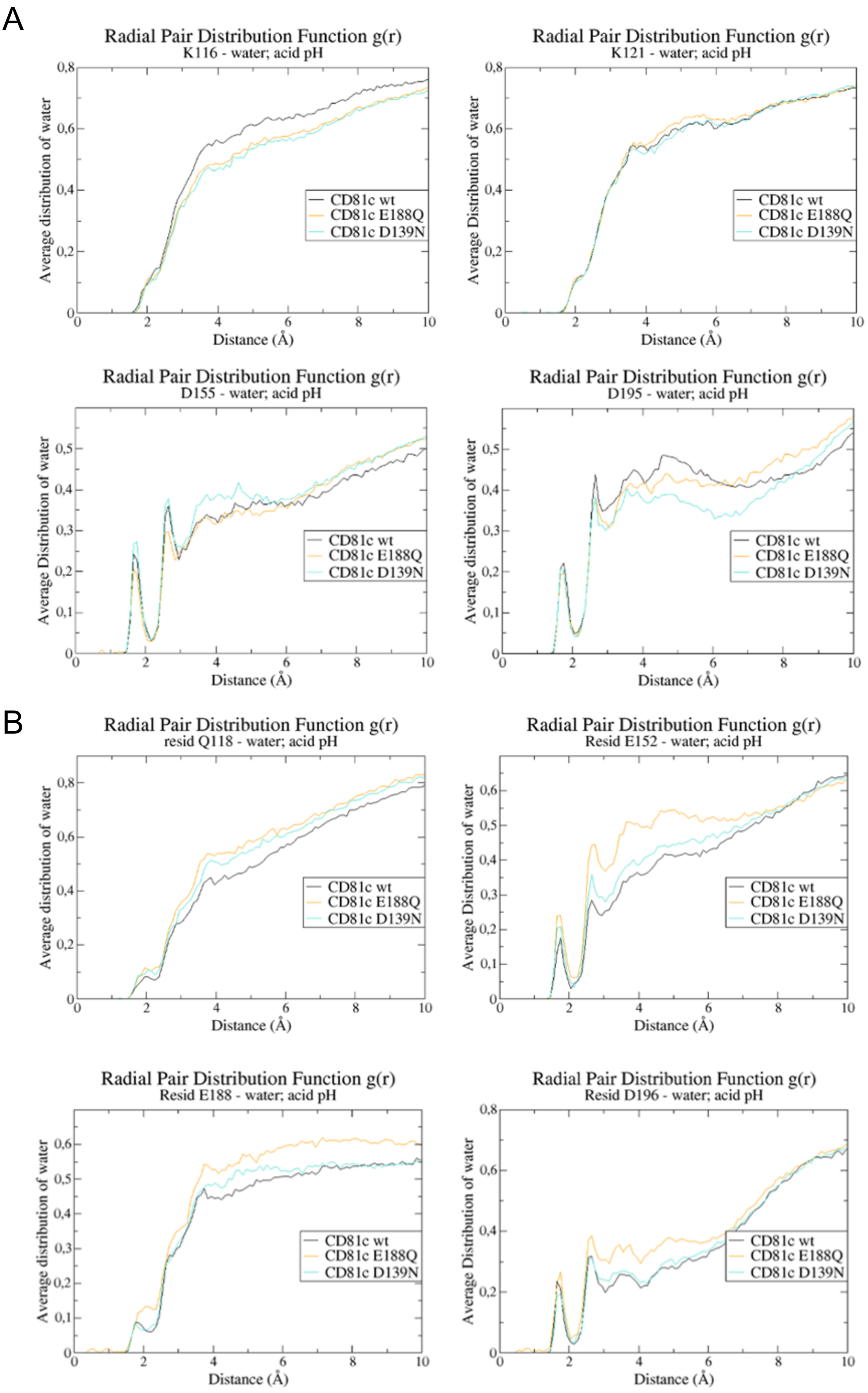
A
Radial Pair Distribution Function g(r)
K116 - water; acid pH
Average distribution of water
Distance (Å)
CD81c wt
CD81c E188Q
CD81c D139N
Radial Pair Distribution Function g(r)
K121 - water; acid pH
Average Distribution of water
Distance (Å)
Radial Pair Distribution Function g(r)
D155 - water; acid pH
Average Distribution of water
Distance (Å)
Radial Pair Distribution Function g(r)
D195 - water; acid pH
Average distribution of water
Distance (Å)
B
Radial Pair Distribution Function g(r)
resid Q118 - water; acid pH
Average distribution of water
Distance (Å)
Radial Pair Distribution Function g(r)
Resid E152 - water; acid pH
Average Distribution of water
Distance (Å)
Radial Pair Distribution Function g(r)
Resid E188 - water; acid pH
Average distribution of water
Distance (Å)
Radial Pair Distribution Function g(r)
Resid D196 - water; acid pH
Average Distribution of water
Distance (Å)

**FIG S2.** (A) Average distribution of water molecules around 10 Å of residues K116, K121, D155 and D195 at low pH (pH 4.6) in $CD81_{LEL}$ close. (B) Average distribution of water molecules around 10 Å of residues Q118, E152, E188 and D196 at low pH (pH 4.6) in $CD81_{LEL}$ close [1].

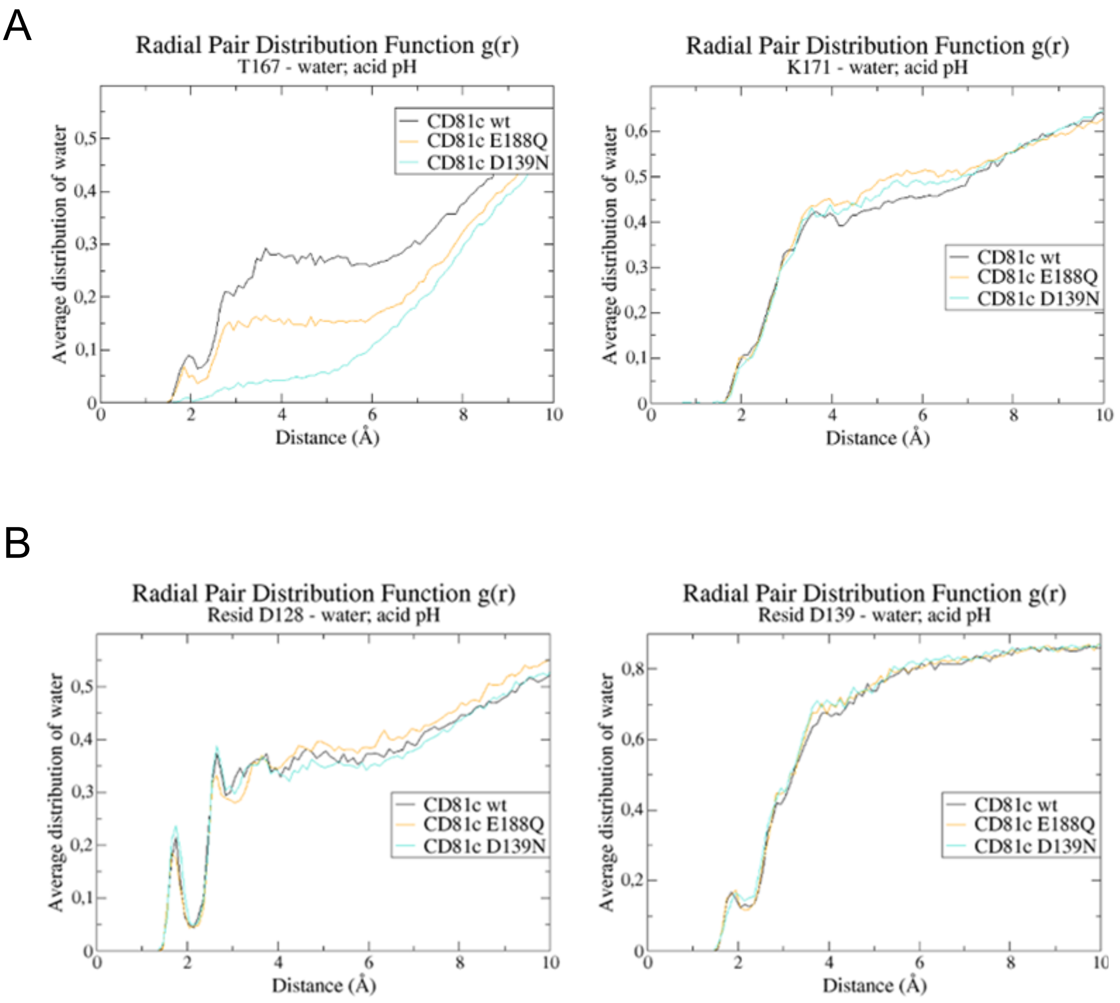


**FIG S3.** (A) Average distribution of water molecules around 10 Å of residues T167 and K171 at low pH (pH 4.6) in $CD81_{LEL}$ close. (B) Average distribution of water molecules around 10 Å of residues D128 and D139 at low pH (pH 4.6) in $CD81_{LEL}$ close [1,2].

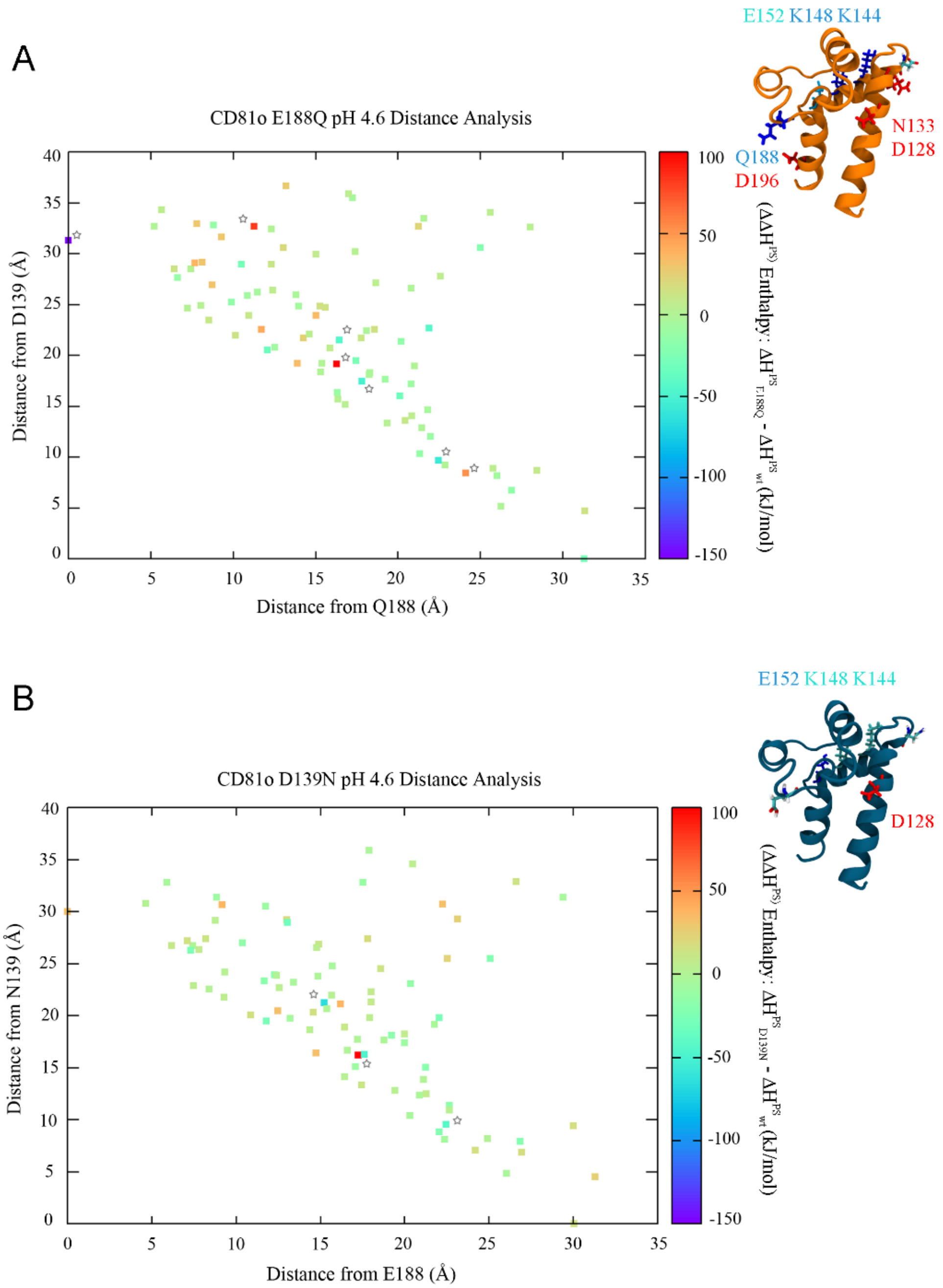


**FIG S4. Distance Analysis on CD81$_{LEL}$ open**. (A) $\Delta\Delta H^{PS}$ of CD81$_{LEL}$ open with mutation E188Q. (B) $\Delta\Delta H^{PS}$ of CD81$_{LEL}$ open with mutation D139N [3].
Stars highlight the position of the residues that get stabilized (blue) and destabilised (red) which are depicted as sticks in the cartoon protein structures.

A

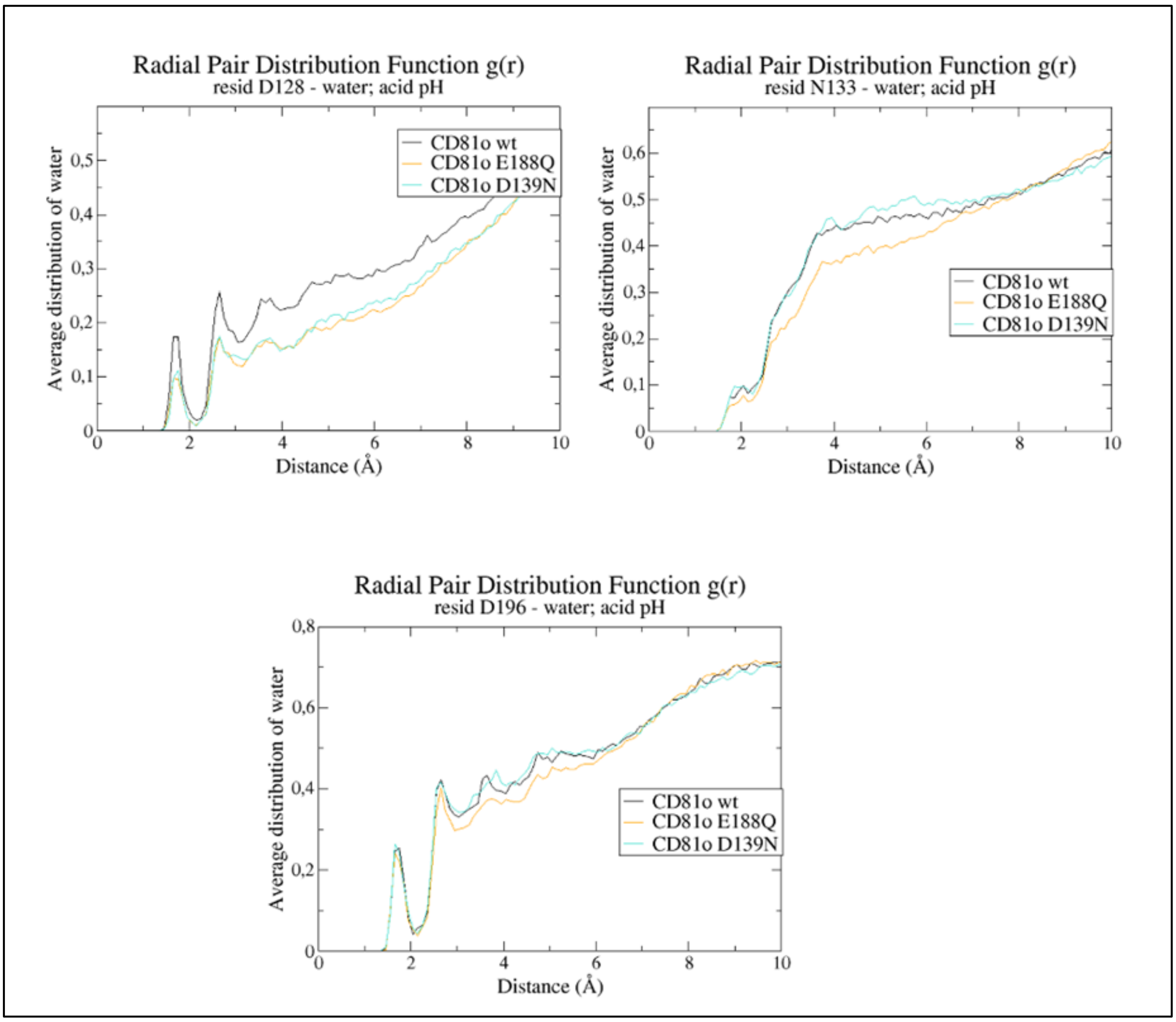
Radial Pair Distribution Function g(r)
resid D128 - water; acid pH
Average distribution of water
Distance (Å)
CD81o wt
CD81o E188Q
CD81o D139N
Radial Pair Distribution Function g(r)
resid N133 - water; acid pH
Average distribution of water
Distance (Å)
CD81o wt
CD81o E188Q
CD81o D139N
Radial Pair Distribution Function g(r)
resid D196 - water; acid pH
Average distribution of water
Distance (Å)
CD81o wt
CD81o E188Q
CD81o D139N

B

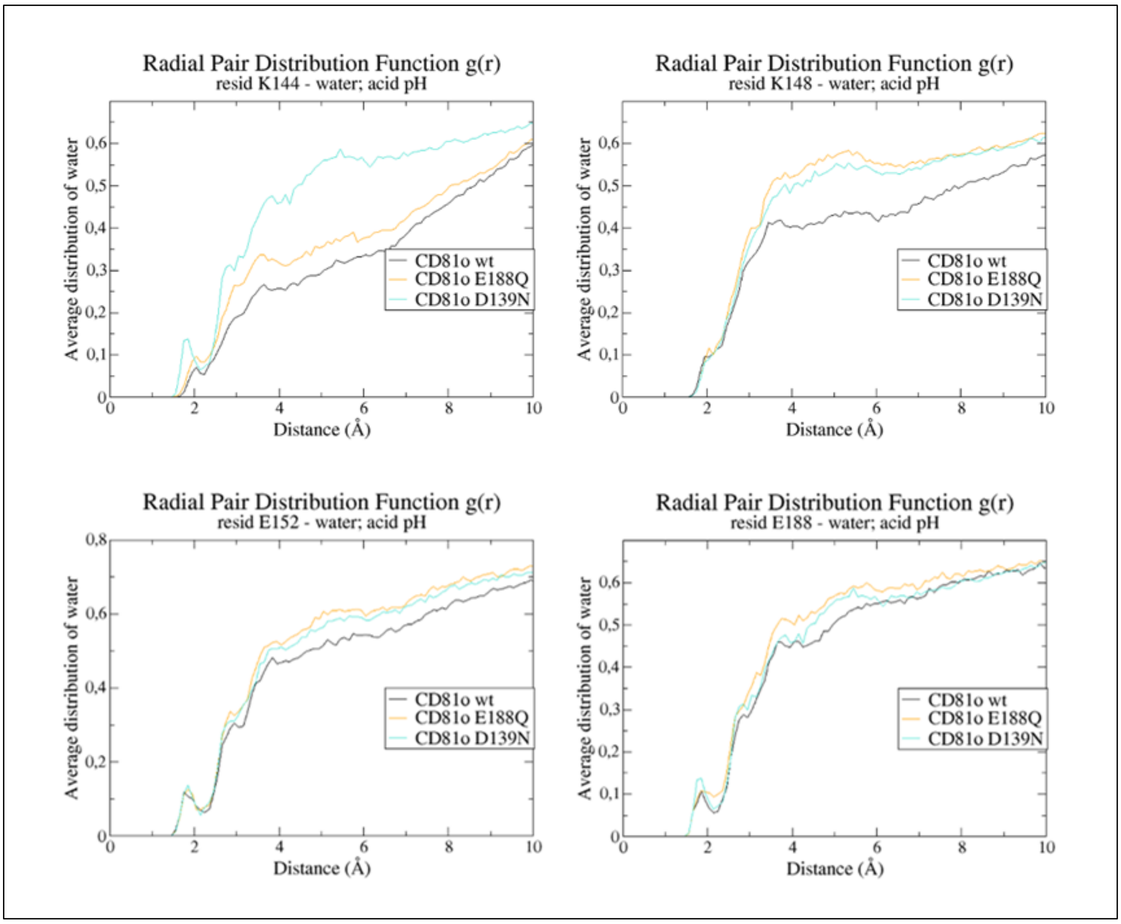


**FIG S5.** (A) Average distribution of water molecules around 10 Å of residues D128, N133, D196 at low pH (pH 4.6) in $CD81_{LEL}$ open. (B) Average distribution of water molecules around 10 Å of residues K144, K148 and E152 at low pH (pH 4.6) in $CD81_{LEL}$ open [1,2].

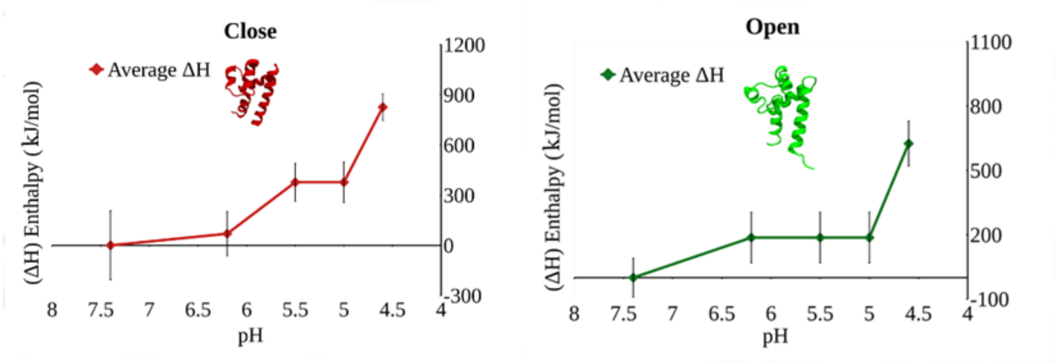


**FIG S6.** Difference of the protein enthalpy at the different endosomal pH between the close and open conformation of $CD81_{LEL}$.

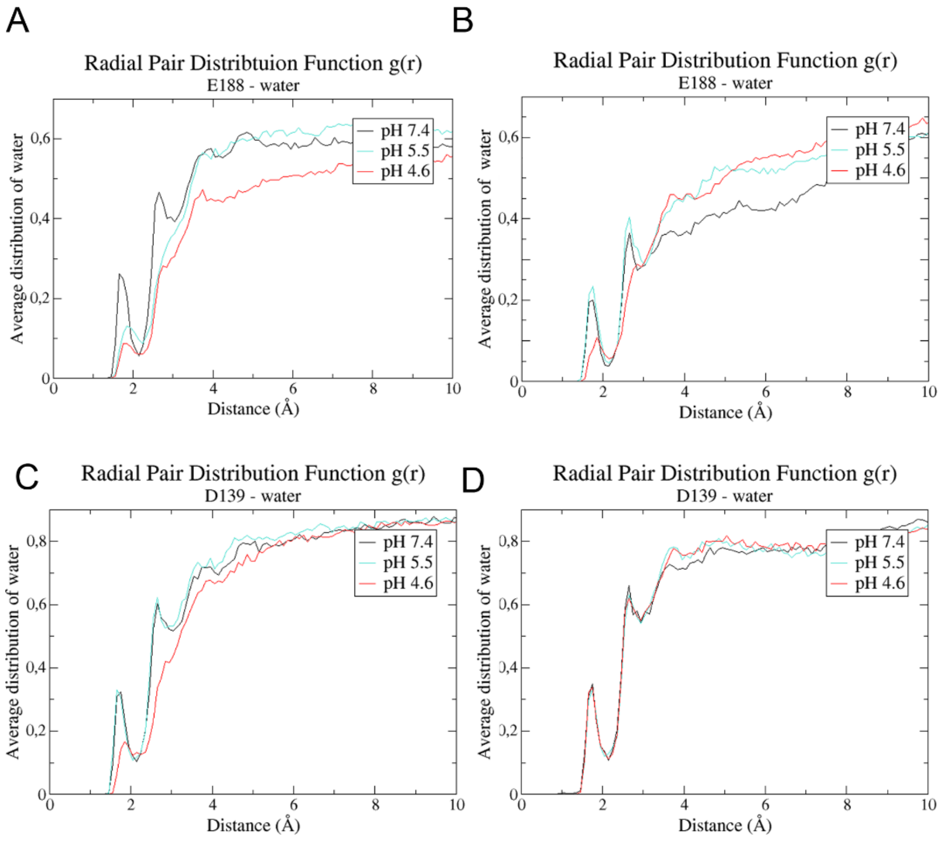


**FIG S7**. **Radial Pair distribution Analysis g(r) on the CD81$_{LEL}$ open-close conformers**. (A-B) Average distribution of water molecules around 10 Å of positions E188 or D139 of CD81$_{LEL}$ close at pH 7.4, 5.5 and 4.6. (C-D) Average water molecules distribution around 10 Å of positions E188 or D139 of CD81$_{LEL}$ open at pH 7.4, 5.5, 4.6 [1,2].

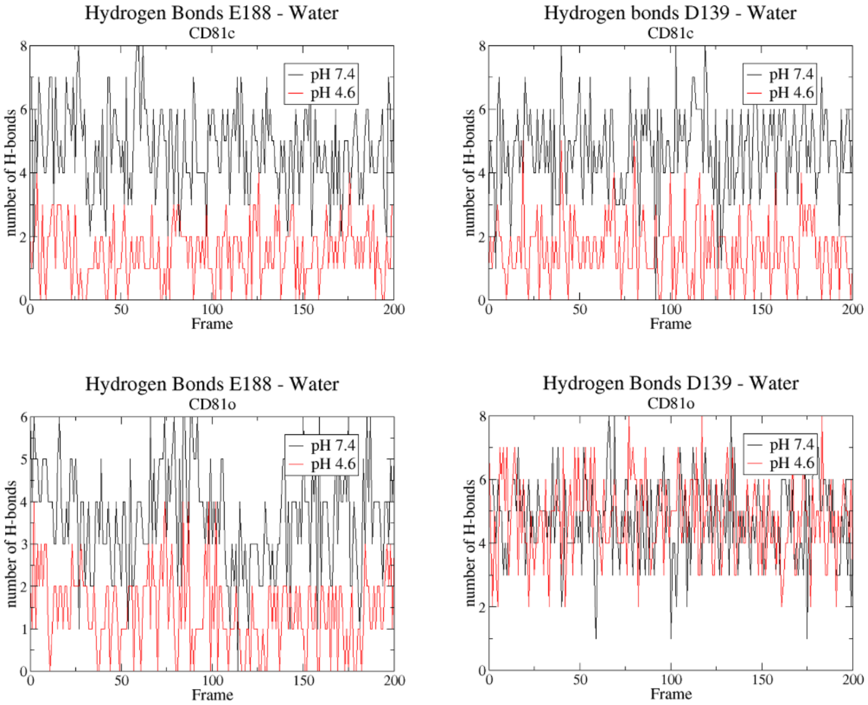


**FIG S8.** Differences in the hydrogen bond network of CD81c at position E188 at pH 7.4 (left) and pH 4.6 (right) within the solvation shell [2].

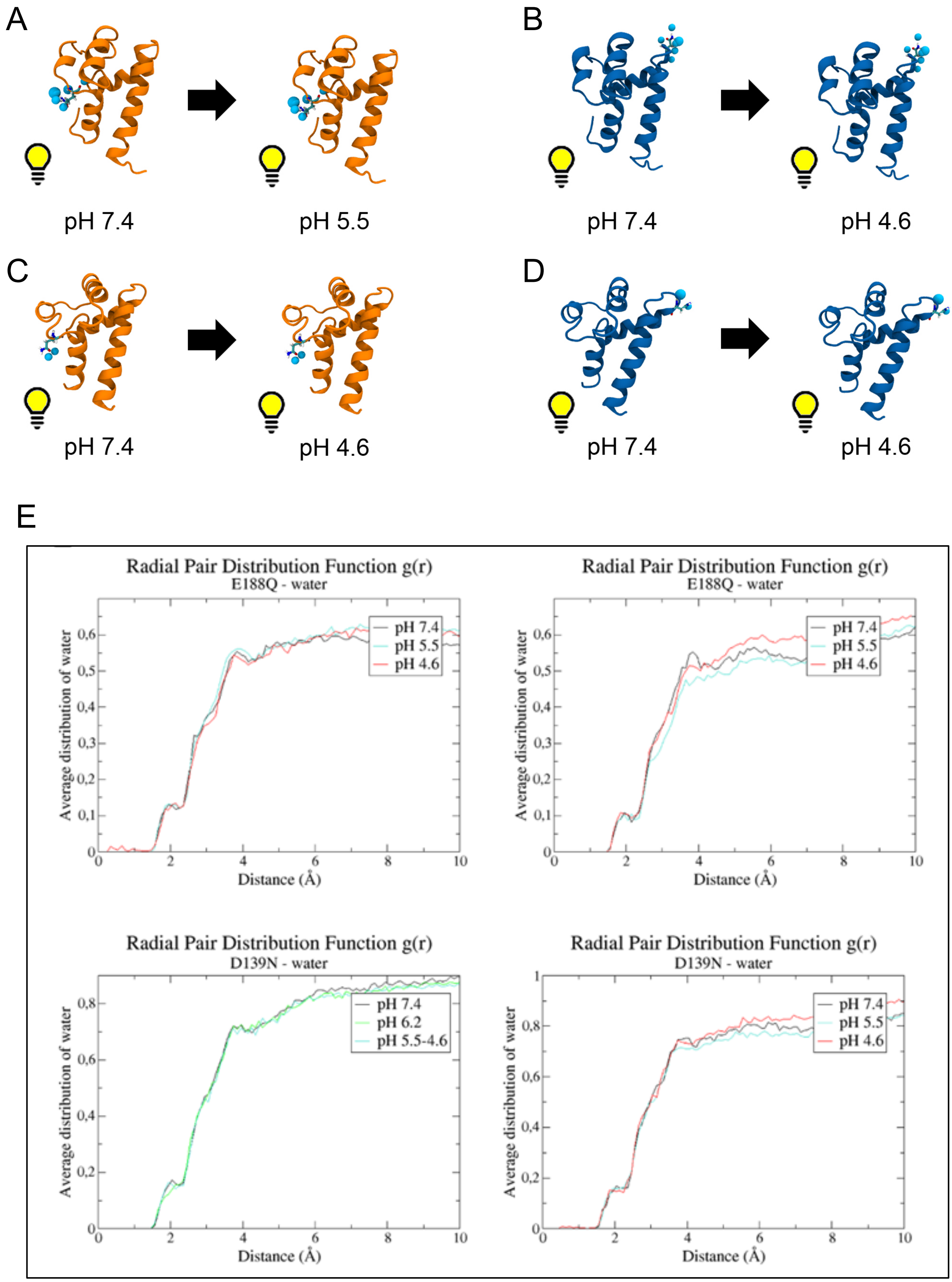


**FIG S9. Mutations D139N and E188Q disrupt the signal for the allosteric transition of the CD81$_{LEL}$ conformers**. (A-B) Residues N139 and Q188 maintain their charge (switched on) at low pH, stabilizing the close conformation in water at low pH 5.5 and 4.6, respectively; (C-D) Similarly in the open conformation, residues N139 and Q188 are not able to sense pH, retaining its level of hydration. Water molecules at 2 Å distance are depicted as cyan spheres. (E) Average distribution of water molecules around 10 Å of positions E188Q or D139N of $CD81_{LEL}$ along endosomal pH [1,2].

A

B

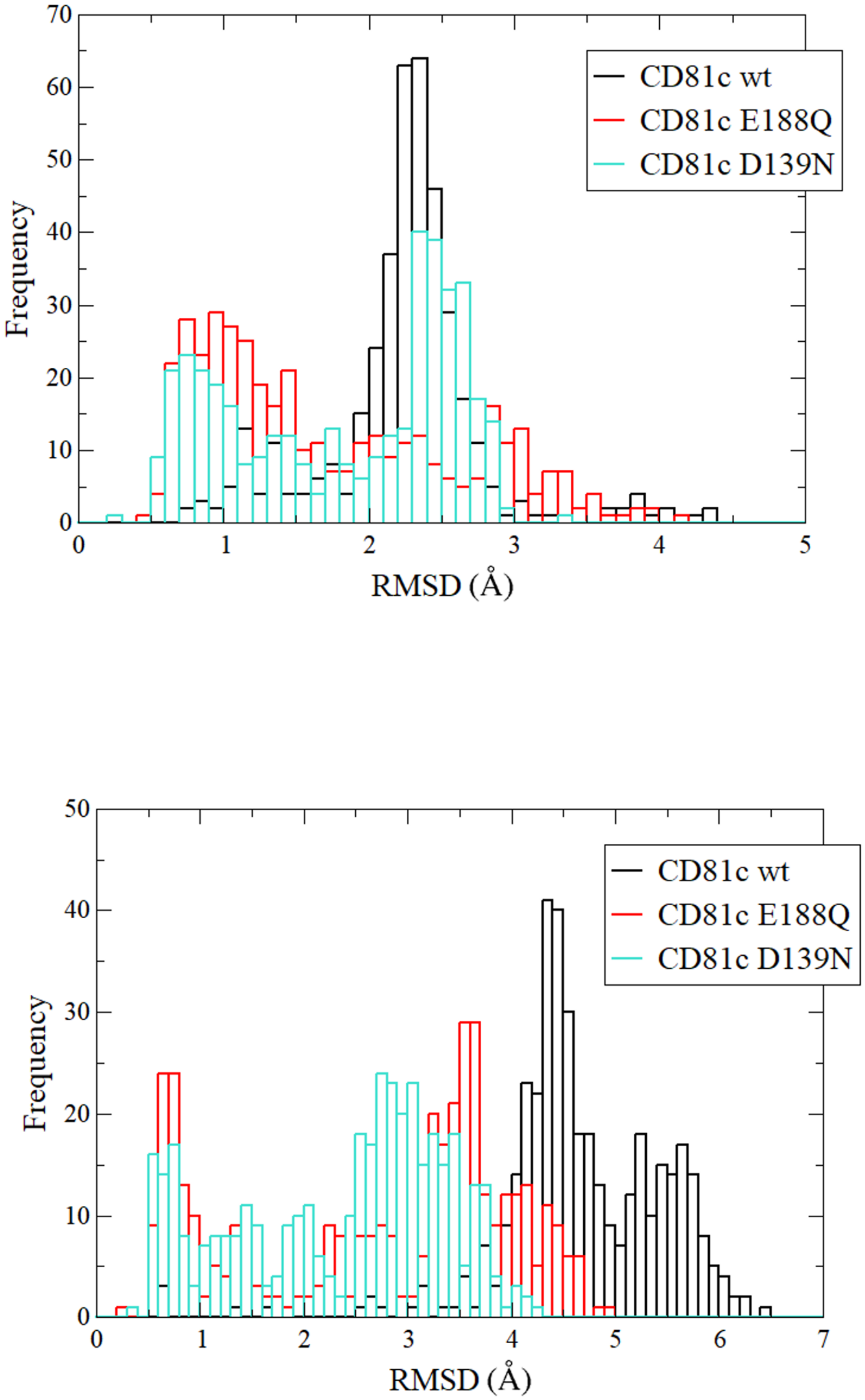


**FIG S10**. Distribution histogram of the effect of mutations D139N and E188Q over the dynamics of $CD81_{LEL}$ head subdomain along 1 µs trajectory at pH 5.5 (A) and pH 4.6 (B) [1].

A

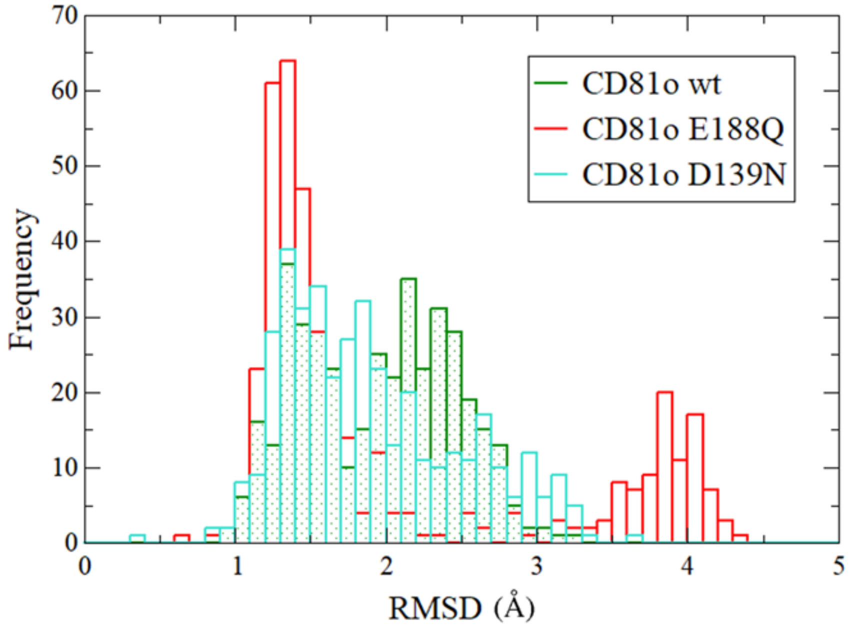


B

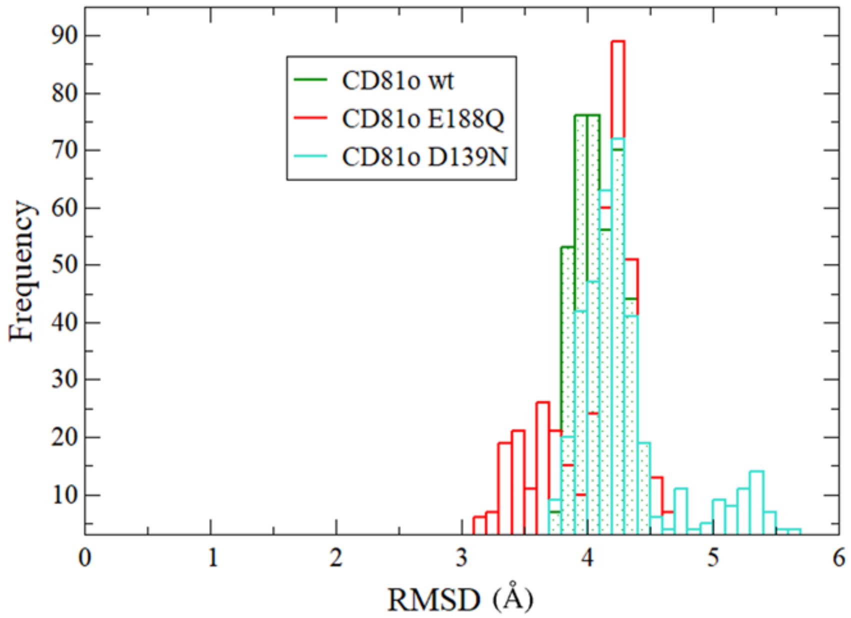


**FIG S11.** Distribution histogram of the effect of mutations D139N and E188Q on the dynamics of $CD81_{LEL}$ open along 1 μs trajectory at (A) pH 4.6 (B) comparing with the reference crystal close structure of $CD81_{LEL}$ (PDB ID 5M3T) [1,4].

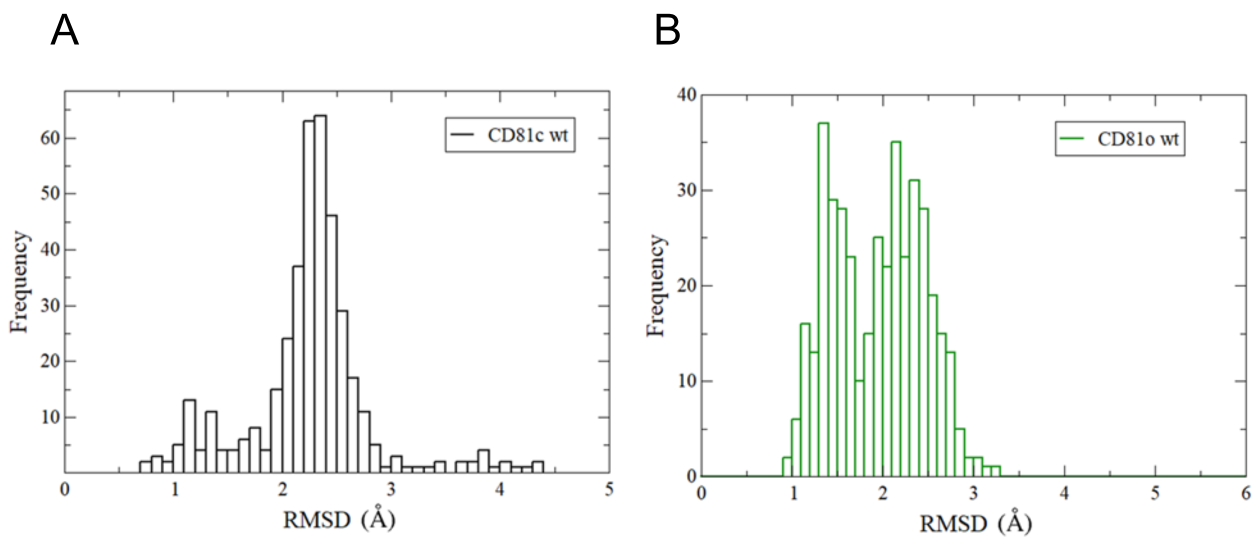


**FIG S12**. Dynamic oscillations explored by CD81$_{LEL}$ head subdomain wild-type in close or open conformation along 1 µs trajectory.

A

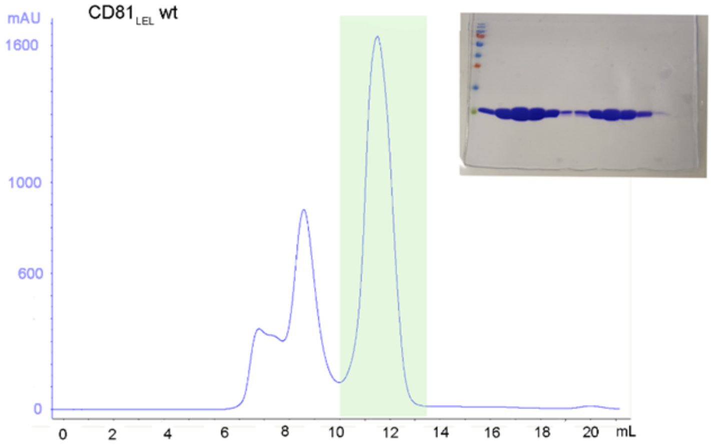
CD81LEL wt
mAU
1600
1000
600
0
0 2 4 6 8 10 12 14 16 18 20 mL

B

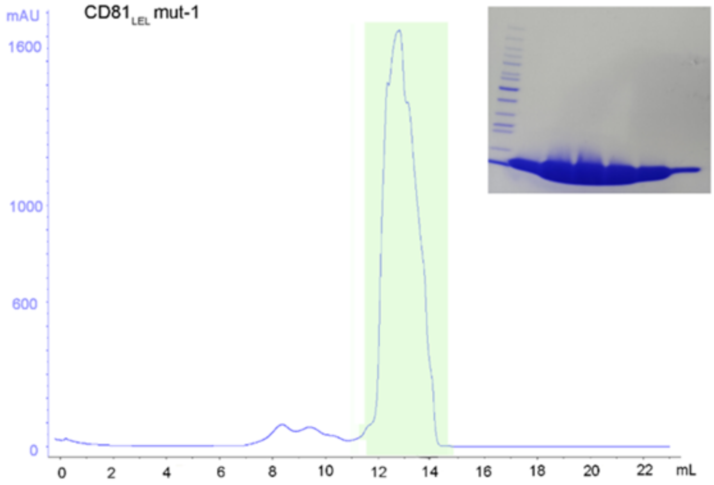
CD81LEL mut-1
mAU
1600
1000
600
0
0 2 4 6 8 10 12 14 16 18 20 22 mL

C

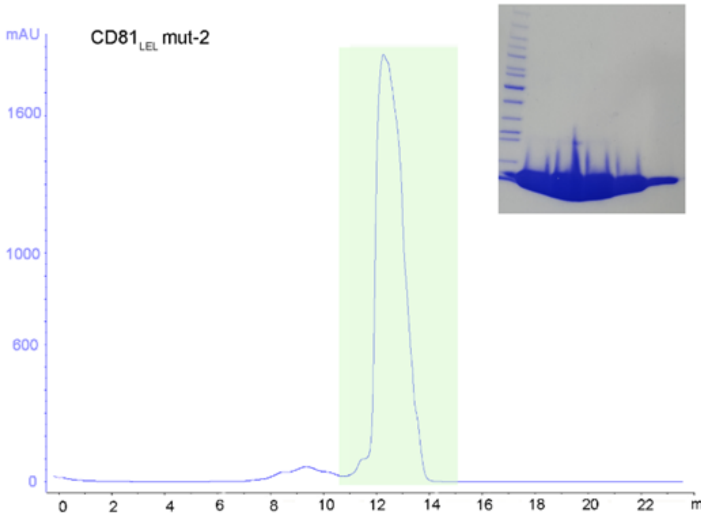
CD81LEL mut-2
mAU
1600
1000
600
0
0 2 4 6 8 10 12 14 16 18 20 22

**FIG S13. Purification of wild-type and mutant $CD81_{LEL}$ proteins.** Elution profile from the size exclusion chromatography of the different $CD81_{LEL}$ proteins; (A) $CD81_{LEL}$ wild-type, (B) $CD81_{LEL}$ mut-1, (C) $CD81_{LEL}$ mut-2. The highlighted peek in green corresponds to the dimeric oligomer. On the side of each profile the corresponding SDS-PAGE gel showing the elution fractions collected from size-exclusion chromatography.

A

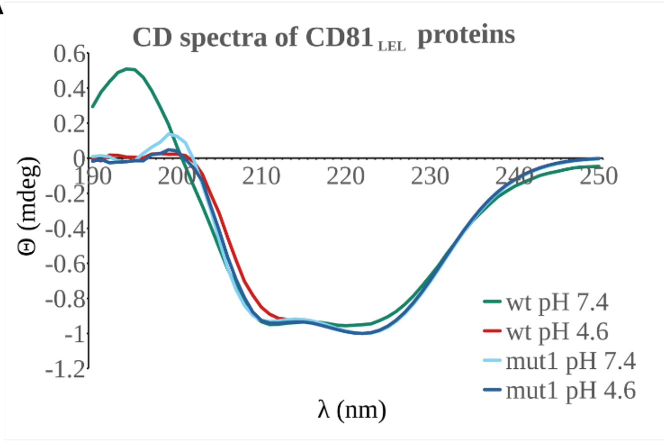


B

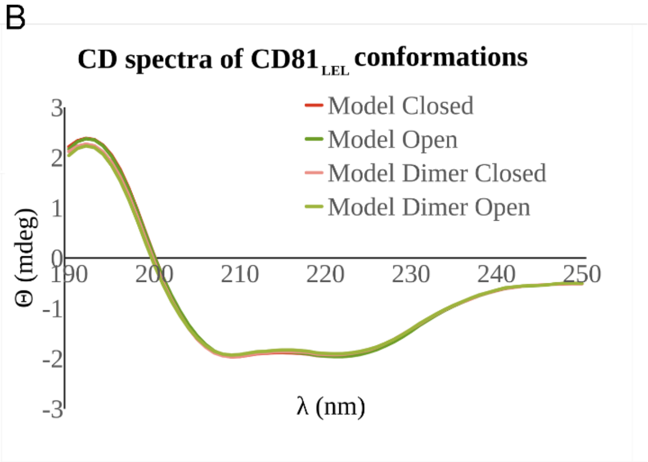


**FIG S14. Secondary structure profile of $CD81_{LEL}$ constructs**. (A) These scans indicate the wild-type and mutant-1 construct display an α-helical structure with little difference between the systems expressed. (B) Comparison of predicted a-helical structure between the open and close structure of $CD81_{LEL}$. As shown, there are imperceptible differences in the CD scan.

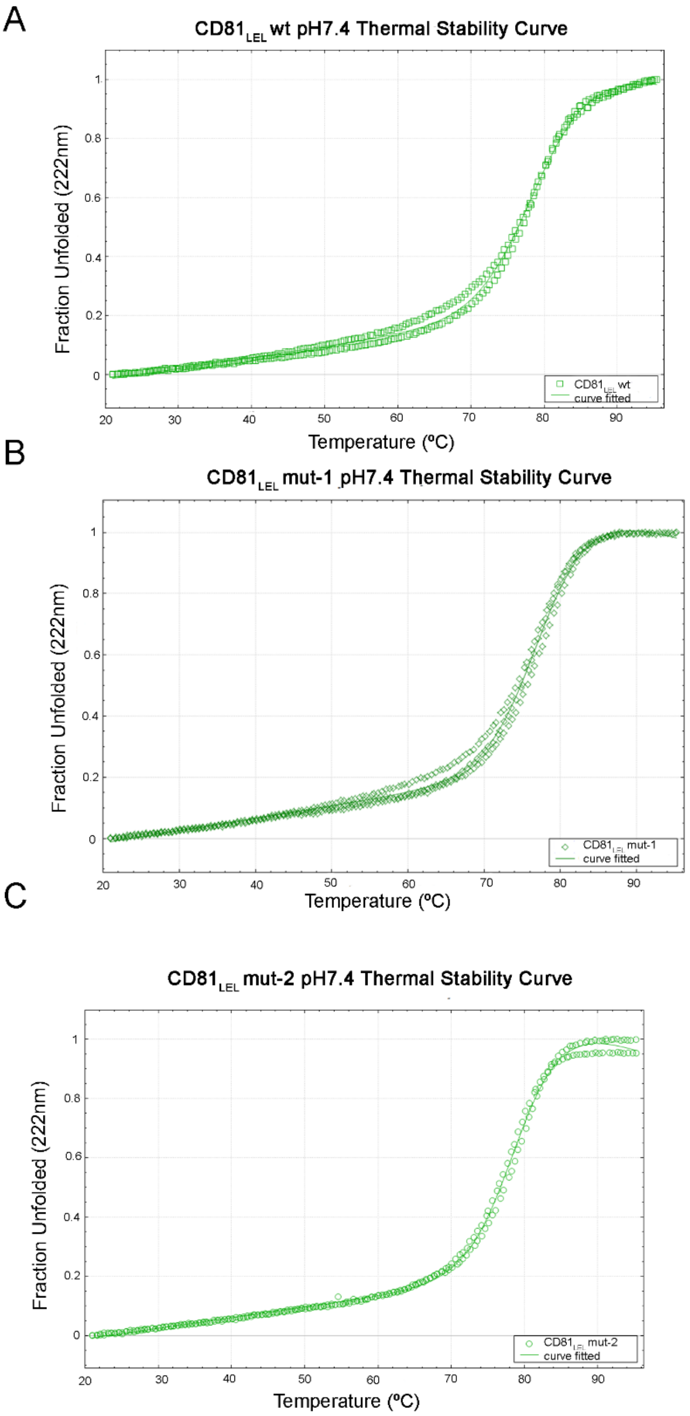


**FIG S15. Thermal stability of the CD81$_{LEL}$ constructs at physiological pH 7.4**. (A) wild-type; (B) Mut-1 (D139N); (C) Mut-2 (E188Q) [5].

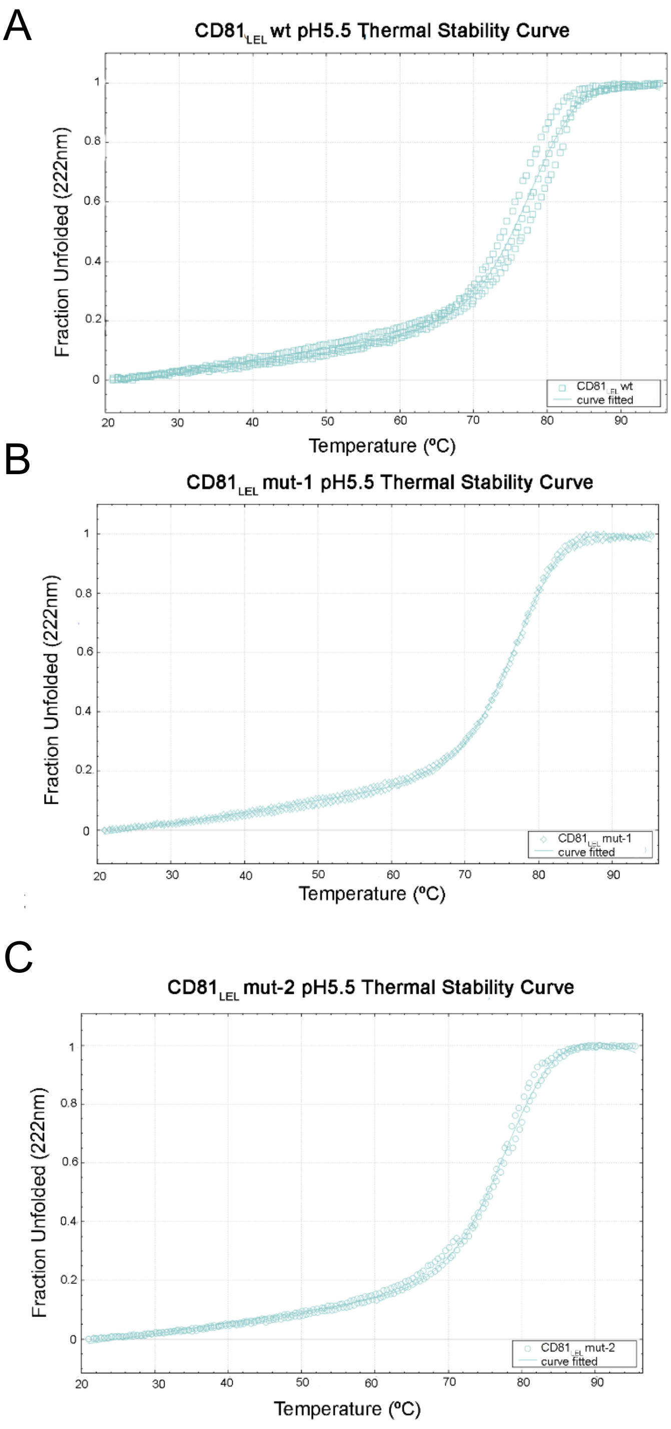


**FIG S16. Thermal stability of the $CD81_{LEL}$ constructs at moderate acid pH 5.5**. (A) wild-type; (B) Mut-1 (D139N); (C) Mut-2 (E188Q) [5]

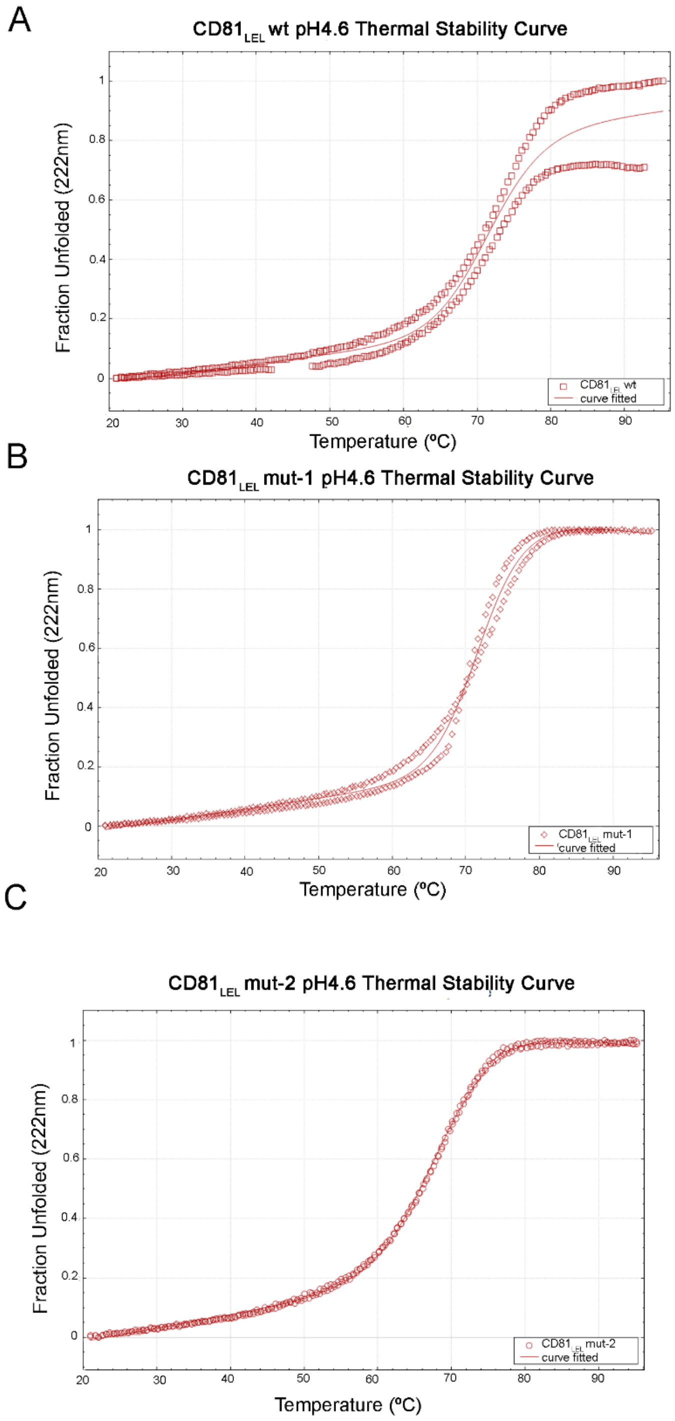


**Supplementary Figure 17. Thermal stability of the $CD81_{LEL}$ constructs at low acid pH 4.6**. (A) wild-type; (B) Mut-1 (D139N); (C) Mut-2 (E188Q) [5].
